# Deletion of Ferritin Heavy Chain Limits Tumor Growth and Promotes Iron-Dependent Stress in Medulloblastoma

**DOI:** 10.64898/2026.07.23.739635

**Authors:** F. Segui, J. Durivault, M. Pagnuzzi, V. Vial, A. Bernini, D. Filipponi, T. Harayama, P. Perné, D. Debayle, K. Muller, E. Pasquier, M. Le Grand, Parks SK., Y. Cormerais, J. Pouyssegur, M. Vucetic, V. Picco

## Abstract

Iron is essential for tumor proliferation and metabolic adaptation but becomes cytotoxic when unbuffered, creating a potential metabolic vulnerability. Ferritin, a conserved iron-storage complex, limits labile iron and establishes the upper threshold of iron tolerance in cancer cells. Here, we report the first ferritin heavy chain (FTH) knockout in a brain tumor model system. Although FTH loss was tolerated under basal conditions through adaptive remodeling of iron metabolism, it exposed profound vulnerabilities under iron stress. FTH deficiency lowered the threshold for iron toxicity, sensitizing medulloblastoma (MB) cells to both canonical ferroptosis and a mechanistically distinct iron-dependent cell death pathway. Oxidative iron stress impaired tumor growth and prolonged survival in orthotopic xenografts, whereas vitamin C–induced iron reduction triggered a selective, iron-dependent, but non-ferroptotic elimination of MB-like cells in tumor organoids. Notably, sensitivity to iron toxicity correlated strongly with cellular phenotype, with mesenchymal-like cells displaying greater susceptibility than epithelial-like counterparts. Collectively, these findings identify ferritin as a central regulator of iron tolerance in MB and establish iron toxicity, not via iron deprivation, as a therapeutically exploitable vulnerability. More broadly, this work provides a mechanistic framework for targeting iron metabolism through modulation of ferritin-dependent iron buffering and iron redox homeostasis in cancers.

**Graphical abstract:** 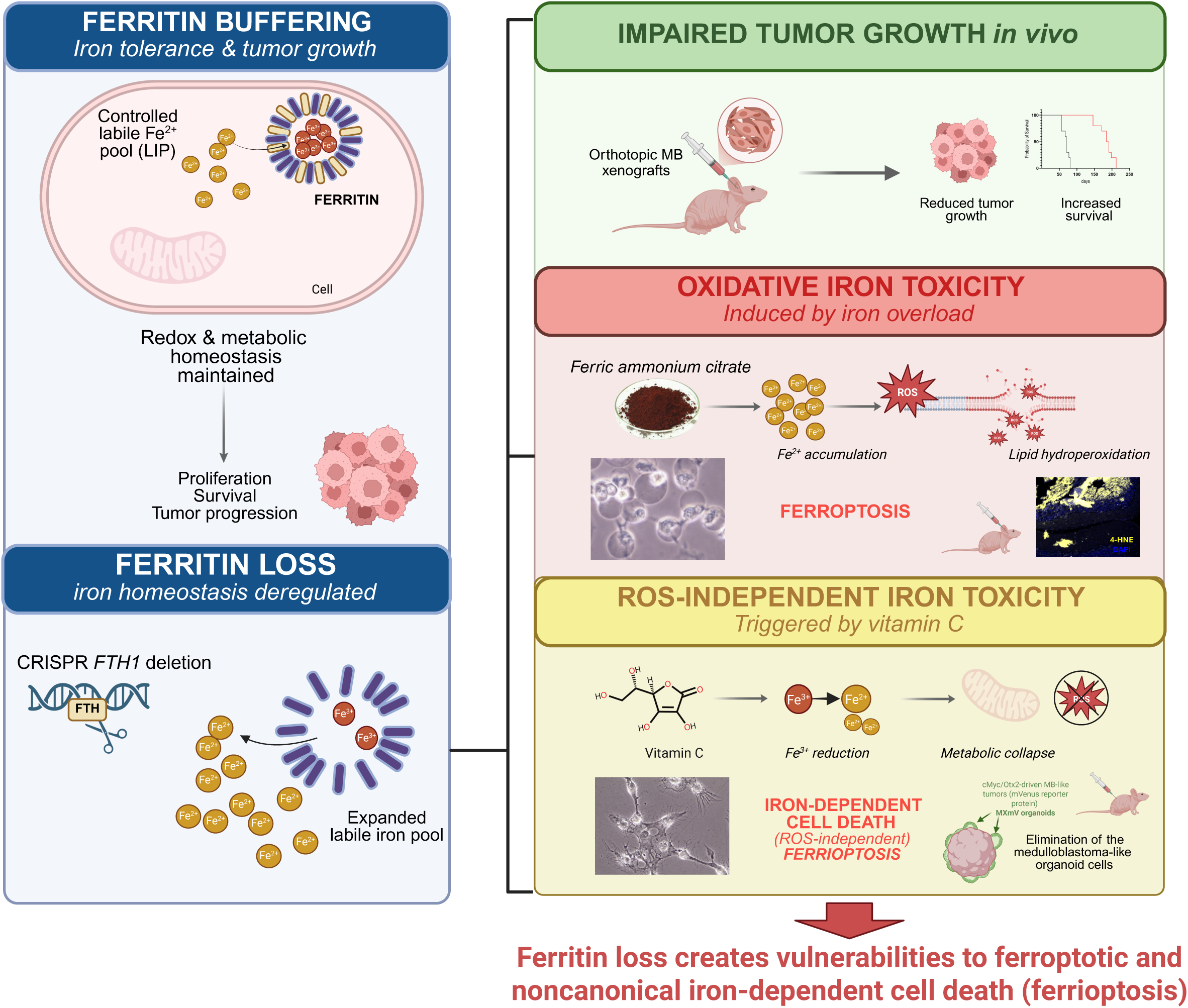

## Introduction

Life’s dependence on iron developed billions of years ago, when oceans teemed with soluble Fe²⁺ under oxygen-free conditions (1). Early organisms harnessed iron for essential metabolic pathways, exploiting its ability to cycle between Fe²⁺ and Fe³⁺ to drive redox reactions. This legacy has remained embedded across all domains of life. The subsequent rise of atmospheric oxygen dramatically altered this landscape, precipitating iron as insoluble oxides and causing a sharp reduction in its bioavailability. In response, organisms evolved sophisticated systems for iron acquisition, storage, and regulation to ensure a reliable supply for critical processes, including DNA synthesis, mitochondrial respiration, and redox metabolism, even in iron-starved environments.

Cancer cells exploit these primordial iron-management programs to meet the demands of accelerated metabolism and proliferation. Tumors upregulate iron uptake, enhance storage, and rewire regulatory pathways to sustain a steady supply of this essential cofactor (2–5). Furthermore, poorly differentiated cancer cell populations, often referred to as cancer stem cells (CSCs), exhibit more pronounced iron dependence reflecting their high metabolic activity and self-renewal capacity (6–9). Yet, iron is a double-edged sword because excess labile iron catalyzes Fenton chemistry, generating reactive oxygen species (ROS), that threaten DNA, protein, and membrane integrity (10,11). To counteract this toxicity, cells rely on ferritin, a highly conserved, multimeric protein capable of sequestering thousands of iron atoms in a redox-inert Fe³⁺ form. Ferritin is a 24-subunit spherical nanocage composed of heavy (FTH) and light (FTL) chains. The FTH subunit possesses ferroxidase activity that catalyzes the oxidation of ferrous (Fe²⁺) to ferric (Fe³⁺) iron, enabling the safe sequestration and mineralization of iron within its hollow core. By buffering cytosolic iron, ferritin regulates iron-dependent signaling and limits oxidative stress (6,7,12–19). Ferritin overexpression is common in cancer, particularly in aggressive and poorly differentiated cells, enabling them to tolerate high iron loads while sustaining proliferation under metabolic stress (2–4,6).

Broad targeting of iron metabolism in cancers through chelation or uptake inhibition has been studied extensively. However, specifically targeting ferritin represents an underexploited vulnerability. Directly perturbing ferritin function could deprive cancer cells of a critical iron reservoir revealing their vulnerability to changing iron concentration and/or oxidative stress, thus selectively compromising their survival. Medulloblastoma (MB), the most common malignant pediatric brain tumor, presents an ideal model for exploring such interventions. High-risk MB subgroups, harbor populations of undifferentiated cells and present poor survival outcomes, despite surgery, craniospinal irradiation, and intensive chemotherapy (20,21). This underscores the urgent need for biologically targeted strategies that exploit unique metabolic vulnerabilities in MB and other cancers.

Here, we investigate the role of ferritin in regulating iron tolerance in MB and assess whether iron redox manipulation can be harnessed therapeutically. Using genetic deletion models, and pharmacological iron modulation we show that ferritin is dispensable for basal survival but essential for protecting MB cells from an iron overload–induced redox catastrophe. Our results were further consolidated using, transformed brain organoids MB models and orthotopic tumor xenografts., We further show that high dose of vitamin C (VitC) induces rapid, iron-dependent, but ROS-independent and non-ferroptotic cell death and selectively eradicates MB-like tumors. Finally, we explore tumor cell states (epithelial vs mesenchymal) as potential key determinants of iron sensitivity.

Collectively, our findings establish that iron toxicity, rather than iron deprivation, is a potent therapeutic vulnerability in MB. We reveal the ferritin buffering capacity acts as a central regulator of tumor survival under stress. By linking iron biology to MB cell metabolism, and ferritin function, this work provides a mechanistic framework for developing iron-based metabolic therapies against aggressive pediatric brain tumors with translational relevance for other solid tumors.

## Material and Methods

### Cell Lines and Culture Conditions

Human MB DAOY and HD-MB03 cells were obtained from the American Type Cancer Collection (ATCC, Manassas, VA, USA) and authenticated in 2026 (Eurofins Genomics, France). They were routinely tested for Mycoplasma (PlasmoTest Mycoplasma Detection Kit; InvivoGen) and cultivated up to a 10^th^ passage. Cells were grown at 37 °C/5% CO_2_ in DMEM (Gibco, Thermo Fisher Scientific Inc, MA, USA) supplemented with 8% Gibco™ Fetal Bovine Serum (FBS, certified, heat inactivated, Thermo Fisher Scientific Inc, MA, USA). SLC7A11 (xCT) deleted cell lines (xCT KO, (22)) were maintained, and experiments were conducted in the same media supplemented with 1 mM N-acetylcysteine (NAC; Sigma-Aldrich (Merck), St. Louis, MO, USA) or 1 μM vitamin E (α-tocopherol; Sigma-Aldrich (Merck), St. Louis, MO, USA). The chemicals and fluorescent probes used in the study are listed in the Table 1 and 2.

**Table 1.** Compounds used in the study.

| Compounds | Function | References |
| --- | --- | --- |
| Ferric Ammonium Citrate (FAC) | Donor of iron ($\text{Fe}^{3+}$) | Sigma 1185-57-5 |
| Vitamin C (VitC) | Iron reducer | Sigma A5960 |
| Deferoxamine (DFO) | Iron chelator | Sigma D9533 |
| Ferrostatin-1 (Fer-1) | Ferroptosis inhibitor | Sigma SML0583 |
| Necrostatin-1 (Nec-1) | Necroptosis inhibitor | ab141053 |
| Pan-caspase inhibitor (Q-VD-OPh) | Apoptosis inhibitor | TargetMol T0282 |
| N-acetylcysteine (NAC) | Antioxidant | Sigma A9165 |
| Basic Fibroblast Growth Factor (bFGF) | Growth factor | StemCell Tech 78003.2 |
| Bafilomycin A1 | Autophagy inhibitor | StressMarq Biosci ST-SIH-391 |

### Genetic deletion of ferritin heavy chain (FTH) using CRISPR-Cas9

DAOY and HD-MB03 wildtype (WT) cells were transfected using jetPEI (Polyplus, 101000053) according to the manufacturer’s instructions with pSpCas9(BB)-2A-GFP (PX458) plasmid (a gift from Feng Zhang; Addgene plasmid #48138; http://n2t.net/addgene:48138; research re-source identifier: Addgene_48138) (23) containing clustered regularly interspaced short palindromic repeats (CRISPR)-CRISPR-associated protein 9 (Cas9) targeting the following regions: *FTH1* exon 1 immediate flanking region gRNA (5′-**G**TTACCTGTCCATGgtgagcg - 3′), and *FTH1* exon 3 gRNA (5′- gggttccaatactcacATGG - 3′). Bold **G** in 5’ is added due to the transcription initiation requirement of a ‘G’ base for human U6 promoter. GFP-positive cells were single-cell sorted by flow cytometry 24 h post-transfection into 96-well plates containing DMEM supplemented with 8% FBS. Knockout clones were screened by immunoblotting and validated by Sanger sequencing (Eurofins Genomics, France). Two independent knockout clones of each cell lines were selected for experiments, to minimize clonal effects.

### Proliferation assay

DAOY and HD-MB03 WT/FTH KO cells were seeded in 6 well plates. Cell proliferation was assessed by trypsinization and counting (Coulter Z1; Beckman Coulter, Brea, CA, USA) after 1, 3 and 7 days. The cell proliferation index was calculated as the percentage of day 1 by standardizing each measurement to the cell number obtained 24 h after seeding (day 1).

### Immunoblotting

Cells lysis was performed in 1.5×Laemmli buffer, and protein concentrations were quantified with the Pierce BCA Protein Assay from (Thermo Fisher Scientific Inc, MA, USA). Protein extracts (15 µg) were subjected to electrophoresis on either 10% or 12% sodium dodecyl-sulfate-polyacrylamide (SDS) gels and subsequently transferred to polyvinylidene difluoride membranes (PVDF, Merck Millipore, Burlington, MA, USA). The membranes were blocked with 5% milk in phosphate-buffered saline (PBS) and incubated with anti-human primary antibodies (Table 3), followed by incubation with the corresponding horseradish peroxidase (HRP)-conjugated secondary antibodies. Immunoreactive bands were detected with horseradish peroxidase anti-mouse or anti-rabbit antibodies (Promega, Madison, WI, USA) using the enhanced chemiluminescence (ECL) system (Merck Millipore, Burlington, MA, USA). Immunoblot analysis was performed using the Li-COR Odyssey Imaging System (Lincoln, NE, USA) and Fusion Fx7 absolute (Vilber, Collégien, France).

### Colony Formation Assay

DAOY and HD-MB03 WT and FTH KO cells were seeded with DMEM + 8% FBS into 60 mm dishes with initial cell numbers of 1000 and 2000 respectively. Cells were treated the next day with different concentration of ferric ammonium citrate (FAC, Thermo Fisher Scientific Inc, MA, USA), erastin (Sigma-Aldrich (Merck), St. Louis, MO, USA) supplemented or not with ferrostatin-1 (Fer-1, Sigma-Aldrich (Merck), St. Louis, MO, USA), as a ferroptosis inhibitor, or DFO (Sigma-Aldrich (Merck), St. Louis, MO, USA), as a known iron chelator. Approximately 7-8 days after treatment, when visible colonies had formed, dishes were stained with 5% Giemsa (Fluka (Sigma-Aldrich), Switzerland) for 30 – 45 min to visualize colonies.

### Flow Cytometry

A total of 10,000 events per sample were analyzed using a BD FACSMelody cytometer (Becton Dickinson (BD) Biosciences, NJ, USA), and data was processed using the FlowJo software version vX.0.7 (Ashland, OR, USA). Cells were cultured in 6-well plates, in triplicate for each condition, and maintained at 37 °C in a 5% CO_2_ atmosphere with their respective media.

***Cell death*** was assessed using propidium iodide (Table 2) (PI; Invitrogen, Thermo Fisher Scientific Inc, MA, USA). DAOY cell lines were seeded at a density of 150 000 cells/well whereas HD-MB03 were seeded at a density of 50 000 cells/well. Both floating and adherent cells were collected, centrifuged, and then resuspended in FACS buffer (PBS, 0.2% bovine serum albumin (BSA), and 2 mM EDTA), with 2 µg/mL of PI added just before analysis.

**Table 2.** Chemical probes used in the study.

| Chemical probes | Function | References |
| --- | --- | --- |
| Propidium iodide (PI) | Cell death measurement | Invitrogen 2015554 |
| $\text{H}_2\text{DCFDA}$ | ROS detection | Invitrogen 2155525 |
| BODIPY C11 | Lipids hydroperoxides detection | Invitrogen 2970996 |
| FerroOrange | Ferrous iron detection ($\text{Fe}^{2+}$) | Dojindo F374 |

***Lipid hydroperoxides*** content was measured using the BODIPY 581/591 C11 dye (Table 2) (Molecular Probes, Eugene, OR, USA). BODIPY 581/591 C11 dye was added in the media to the final concentration of 2 μmol/L and cells were incubated for 30 minutes at 37 °C/5% CO_2_ protected from the light. Subsequently, cells were washed 2 times with PBS, detached using accutase (Dutscher, Brumath, France), and resuspended in FACS buffer. For data presentation the modal scaling option was used (each peak is normalized to its mode, i.e., to % of maximal number of cells found in a particular bin).

***ROS content*** was measured by using H_2_DCFDA (Table 2) (Abcam, Cambridge, UK). Before adding the probe, cells were washed twice and detached using trypsin. Cells were resuspended in FACS buffer with the H_2_DCFDA probe (2 µM final) for 30 min at 37 °C in a 5% CO_2_ environment, protected from light. Following, FACS analysis of total ROS level was performed, and the data are represented in modal scaling (each peak is normalized to its modal, i.e., to % of the maximal number of cells found in a particular bin).

### Intracellular Fe^2+^ measurement

Ferrous iron was measured using FerroOrange (Table 2) (Dojindo Molecular Technologies, Kumamoto, Japan). After the experiment, cells were washed twice with PBS and resuspended in DMEM media without iron (Thermo Fisher Scientific Inc, MA, USA) suppemented with the FerroOrange probe (1 µM) for 30 min at 37 °C in a 5% CO_2_ environment, protected from light. Fluorescence intensity has been measured with a Xenius UVMC specrfluorimeter (SAFAS, Monaco). Fluorescence is normalized based on cell numbers of each condition.

### Metabolic flux analysis

Oxygen consumption rates (OCR) and extracellular acidification rates (ECAR) were measured using the Seahorse XF HS Mini extracellular flux analyzer (Agilent Seahorse Bioscience, North Billerica, MA, USA). Cells were seeded in XF-compatible Seahorse plates and cultured for 24 h to allow formation of a confluent monolayer. Prior to the assay, cells were left untreated or pretreated for 1 h with 10 mM VitC (pH adjusted to 7.2-7.4). One hour before the measurement, the culture medium was replaced with Seahorse assay medium (Agilent Seahorse Bioscience, North Billerica, MA, USA) lacking glucose, pyruvate, serum, and bicarbonate. Plates were incubated in a non-CO₂ incubator at 37 °C for equilibration. The treatment conditions were maintained throughout the entire assay. Basal OCR and ECAR values were recorded, followed by a mitochondrial stress test performed using sequential injections of 1) assay medium or 10 mM VitC (for internal control condition), 1 mM glucose, 3 µM carbonyl cyanide-p-trifluoromethoxyphenylhydrazone (FCCP), and 1 µM rotenone/1 µM antimycin A. Normalization to cell count was performed after each experiment, and data were presented as milli-pH units (mpH) per minute for ECAR and as picomoles of O_2_ per minute for OCR.

## Immunofluorescence

### Cryosections

Tumor samples were recovered from the animals and embedded in the OCT compound according to the manufacturer’s protocol (Thermo Fisher Scientific Inc., Waltham, MA, USA). 5 μm thin sections were prepared with a cryostat (Leica Microsystems, Wetzlar, Germany) fixed in 4% paraformaldehyde (PFA) for 10 min at 4 °C and blocked in 1% horse serum in tris-buffered saline (TBS) for 1 h. The sections were incubated with primary antibody (shown in Table 4) in blocking solution (1% BSA) overnight at 4 °C and thereafter washed with TBS containing 0.1% Tween. Afterward, the preparations were incubated with corresponding fluorescent secondary antibody (Table 4). Next, preparations were washed with TBS containing 0,1% Tween with Hoechst. Fluorescence images of the cells were taken with a DMI4000 (Leica Microsystems, Wetzlar, Germany) inverted microscope equipped with a 10x objective (Leica Microsystems, Wetzlar, Germany) and a Zyla 5.5 camera (Andor Technology Ltd, Belfast, Northern Ireland, UK). Preparations were mounted (Vector Laboratories, Burlingame, CA, USA) and imaged using a Leica microscope (DMI4000B, Leica Microsystems, Wetzlar, Germany).

**Table 3.** Antibodies used for Western blot analysis in the study.

| Antibodies | References |
| --- | --- |
| Ferritin heavy chain (FTH) | CST 4393S |
| Ferritin light chain (FTL) | NBP2-54574 |
| Iron regulatory protein 2 (IRP2) | CST 37135S |
| Transferrin receptor 1 (TFRC1) | Ab 84036 |
| Ferroportin-1 (FPN1) | NBP1 21502 |
| Divalent metal transporter 1 (DMT1) | Ab 55735 |
| Phospho-ribosomal protein S6 kinase (p-p70S6K) | CST 9205 |
| Ribosomal protein S6 kinase p70S6K (p70S6K) | CST 9202S |
| Phospho-ribosomal protein S6 (p-RPS6) | CST 2211 |
| Ribosomal protein S6 (RPS6) | CST 2217S |
| Phospho-eIF4E-binding protein 1 (p-4EBP1) | CST 13443S |
| eIF4E-binding protein 1 (4EBP1) | CST 9452 |
| Microtubule-associated proteins 1A/1B light chain 3B (LC3) | Proteintech 14600-1-AP |
| P62 | Bioscience 610883 |
| Poly(ADP-ribose) polymérase (PARP) | CST 9542S |
| c-Myc | CST 5605 |
| Orthodenticle homeobox 2 (Otx2) | Proteintech 13497-1-AP |
| Neural cadherin (N-cadherin) | CST 13116S |
| Vimentin | CST 5741 |
| Snail | CST 3879S |
| Twist | MAB 6230 |
| Zinc finger E-box binding homeobox 1 (Zeb1) | NBP1 05987 |
| Glutathione peroxidase 4 (Gpx4) | CST 52455 |
| Cystine transporter (xCT) | CST 12691 |
| Acyl-CoA synthetase long chain family member 4 (ACSL4) | Sanat Cruz 271800 |
| Activating transcription factor 4 (ATF4) | CST 11815 |
| Caspase 3 | CST 9662 |
| Beta-Actin | CST D6A8 |
| Beta-Tubulin | CST 2128S |
| ARD1 | Homemade (86) |

**Table 4.** Antibodies used for immunofluorescence staining in the study.

| Antibodies | References |
| --- | --- |
| Nestin | Ab 18102 |
| Ki67 | Ab 16667 |
| 4-Hydroxynonenal (4-HNE) | Ab 46545 |
| Transferrin receptor 1 (TFRC1) | Ab 84036 |

### Paraffin sections

Tumor sections (8 µm) were rehydrated by passing through different xylene ethanol baths at increasing concentrations from 100% to 50%. A demasking step was performed by incubating the slides with sodium citrate (pH 6.0). The sections were then washed twice with TBS + 0.025% Triton x100 and incubated for 30 minutes with TBS + H_2_O_2_. The slides were saturated with 3% BSA and then contacted with primary antibodies (see Table 4) overnight. The slides were rinsed twice and then incubated with corresponding secondary antibodies. Preparations were mounted with mounting medium containing DAPI (Vector Laboratories, Burlingame, CA, USA). Fluorescent images were taken with Zeiss microscope (Carl Zeiss Microscopy GmbH, Jena, Germany) using Zeiss software (Carl Zeiss Microscopy GmbH, Jena, Germany).

### Prussian blue staining

Iron deposits in tumor tissues were detected by Prussian blue staining (Sigma-Aldrich (Merck), St. Louis, MO, USA). Briefly, paraffin-embedded brain tumor sections were deparaffinized, rehydrated, and incubated with a freshly prepared mixture of potassium ferrocyanide and hydrochloric acid, allowing Fe³⁺ to form insoluble Prussian blue complexes. Sections were subsequently counterstained with pararosaniline solution, dehydrated, and mounted for microscopic analysis. Images were taken with Zeiss microscope (Carl Zeiss Microscopy GmbH, Jena, Germany) using Zeiss software (Carl Zeiss Microscopy GmbH, Jena, Germany).

### Drug combination experiment

Briefly, 4,000 living cells were seeded per well with the Certus Flex® (GyGer) in 384-well plates (Corning, #3764) for the drug combination screen. Cells were incubated in the presence of iron homeostasis drugs alone or in association with a single dose of VitC (1µM). Drugs were distributed with an Echo 550 liquid dispenser® (RRID:SCR_027476; Labcyte) at 6 different concentrations covering 3 logs in constant DMSO. Cell viability was measured using CellTiter-Glo® 2.0 Cell Viability Assay (Promega, #G9243) after 72h of drug incubation in a humidified environment at 37°C and 5% CO2. Luminescence was measured using a PHERAstar® plate reader (RRID:SCR_027001; BMG). Data were normalized to negative control wells (DMSO only). IC50, defined as half maximal inhibitory concentration values and AUC (Area Under the Curve – %.mol.L-1) were obtained using library(ic50), library(drc), library(drda), library(ggplot2) and library (PharmacoGx) packages from R studio. Compounds were considered as potentially antagonistic with VitC when the difference in AUC between VitC and CTL conditions is < to -10%.mol.L−1. This cut-off corresponds to more than 3 times the S.D of the AUC of each compound (< 3%) and considered as significant.

### Generation, maintenance and transfection of cerebral organoids, and isolation of TOCs

Human induced pluripotent stem (iPS) cells (SCTi003-A, Stemcell Technologies, Vancouver, Canada) were maintained on a layer of Matrigel™ hESC-qualified (Corning, Thermo Fisher Scientific Inc, MA, USA), in mTeSR Plus medium (Stemcell Technologies, Vancouver, Canada). All cells were mycoplasma free. iPSC were dissociated with ReLeSR, an enzyme-free reagent without manual selection or scraping (Stemcell Technologies, Vancouver, Canada). Organoids were generated with the STEMdiff Cerebral Organoid Kit and the STEMdiff™ Cerebral Organoid Maturation kit (Stemcell Technologies, Vancouver, Canada) based on the protocol published by Lancaster et al. (24,25). At day 35 of differentiation, organoids were electroporated using a nucleofection protocol adapted from Denoth-Lippuner et al. (26). Briefly, five organoids were resuspended in 100 μL of Nucleofector Solution (Cell Line Nucleofector Kit V, Lonza) containing a total of 10 μg of plasmid : 2 μg of hyPBase (PiggyBac hyperactive transposase; pPB[Exp]-EGFP-CMV>hyPBase, VectorBuilder) together with 8 μg of PBCAG-mVenus for the control condition, or 2 μg of hyPBase together with 8 μg of PBCAG-MXmV, expressing c-Myc, OTX2, and mVenus. Electroporation was performed using program A-023 on a Nucleofector 2b device (Amaxa).

After 29 days post-electroporation, organoids were treated with 1 mM of VitC (Sigma-Aldrich (Merck), St. Louis, MO, USA) for 10 days. On day 39 of treatment, VitC concentration was increased to 10 mM and maintained for an additional 10 days. Organoids were harvested on day 49 for western blot analysis. A subset of control organoids received only late high-dose VitC (10 mM) starting on day 49 and continuing until day 63. Tumor development (presence and size) was monitored every 4 days using imaging (Macrofluo, Zeiss, Germany). Upon establishment of an organoid-derived tumor, detached Otx2/c-Myc/mVenus-positive tumor cells were collected and transferred to maturation medium. From this point onward, tumor-organoid cells (TOCs) were maintained as an independent MB-like cell line.

### Intracranial Orthotopic Tumor Xenograft Models

DAOY-Luc (80 000 cells per animal) or HD-MB03-Luc (5 000 cells per animal) were stereotaxically implanted into the cerebellum of 9-week-old Rj: NMRI-Foxn1 nude (nu/nu) female mice (Janvier Labs, Le Genest-Saint-Isle, France). The cells were implanted into the left cerebellar hemisphere (2 mm posterior, 1.5 mm left of the lambda point, and 2.5 mm deep) using a Hamilton syringe fitted with a needle (Hamilton Company, Reno, NV, USA) and following a previously described procedure (27).

### In vivo Fe-dex/VitC treatment

Mice implanted with HD-MB03-Luc WT or FTH KO cells were administered Fe-dex (500 mg/kg; Sigma-Aldrich (Merck), St. Louis, MO, USA) once daily for 5 consecutive days or with VitC (3 g/kg; pH adjusted to 7.2-7.4; Sigma-Aldrich (Merck), St. Louis, MO, USA) two times a day for two weeks *via* intraperitoneal injection, starting 20 days post-implantation. Fe-dex treatment was repeated using the same protocol two weeks later. Treatment initiation was guided by the homogeneous appearance of tumors, as indicated by the luciferase signal. Control mice received PBS on the same schedule. Survival was assessed daily based on neuropathological symptoms, including gait abnormalities and >10% weight loss, which were used as humane endpoints. Each group included at least 10 mice to ensure sufficient statistical power.

### Bioluminescence Imaging

Tumor growth was evaluated by bioluminescent imaging (Newton 7 FT-500, Vilber, Collégien, France) every 2 days after intraperitoneal injection of 3.3 mg of d-Luciferin dissolved in 100 µL of PBS. Ten minutes post-injection, mice were anesthetized with isoflurane and imaged with an IVIS Spectrum. Bioluminescence signals [total flux (photons/second)] were quantified using Kuant software (Newton 7 FT-500, Vilber, Collégien, France).

### Study Approval

All animal experiments were conducted in strict accordance with the recommendations of the Guide for the Care and Use of Laboratory Animals. All animal studies were approved in advance by the local animal care committee (Veterinary Service and Direction of Sanitary and Social Action of Monaco; APAFIS # 19480-2019022616164184v4).

### Lipidomic analysis

Lipids were extracted with the method of Bligh and Dyer. For that, cell pellets were resuspended in 400 µL 50% methanol/water, followed by 300 µL methanol, internal standards (4 µL SPLASH LIPIDOMIX Mass Spec Standard + 2 µL SphingoSPLASH I, all from Avanti Polar Lipids), and 250 µL chloroform. The mixture was shaken for 10 minutes. Then, 250 µL of water and 250 µL of chloroform were added to induce phase separation. The mixture was shaken for 10 minutes, centrifuged for 15 minutes at 3,000 rpm, and 400 µL of the organic phase was transferred to new glass tubes and dried under nitrogen. Dried samples were reconstituted in 60 µL isopropanol:methanol:water (5:3:2). Three µL of extracts were injected into an Ultimate 3000 UHPLC system coupled to a Q Exactive mass spectrometer (Thermo Fisher Scientific).

Liquid chromatogram was done with an Accucore C18 column (150 × 2.1 mm, 2.6 µm particles), using mobile phase A (acetonitrile:water (1:1, v/v) supplemented with 10 mM ammonium formate and 0.1% formic acid) and mobile phase B (isopropanol:acetonitrile:water (88:10:2, v/v) supplemented with 2 mM ammonium formate and 0.02% formic acid) at 400 µL/min. The gradient was: 35% mobile phase B at 0 minutes, 60% B at 4 minutes, 70% B at 8 minutes, 85% B at 16 minutes, 97% B at 25 minutes, 100% B at 25.1 minutes until 31 minutes, and back to 35% B for 4 more minutes.

Mass spectrometry was performed in a data-dependent MS2 mode with top 15 ions being subjected to MS2 analyses, in positive and negative ion modes. MS1 resolution was 70,000, scan range was m/z = 250–1200, AGC target was 1,000,000 and maximum injection time was 250 msec. MS2 analyses were done with isolation windows of m/z = 1, normalized collision energy of 25% and 30% for positive mode and 20%, 30%, and 40% for negative mode, resolution 35,000, AGC target 100,000, and maximum injection time 80 msec.

Peaks were annotated with MS-DIAL 5.5 based on m/z, MS2 spectra, and retention times. Lipid semi-quantification was done by dividing peak areas by those of internal standards and multiplying by the quantities (in pmol) added to the samples.

### Transcriptomic dataset

Transcriptomic data from primary MB samples were obtained from the publicly available GSE85217 cohort (28), generated using the Affymetrix Human Gene 1.1 ST microarray platform. Preprocessed, gene-level normalized expression data were directly retrieved from the Gene Expression Omnibus (GEO) repository. This dataset includes primary tumors representing the four principal molecular subgroups of MB: WNT, SHH, Group 3, and Group 4. The expression matrix was transposed to obtain a sample- by gene format and molecular subgroup annotations were extracted from the metadata and harmonized into four categories (WNT, SHH, Group3, Group4). Samples lacking subgroup annotation were excluded from downstream analyses. The final cohort used from downstream analysis included N = 763 samples, distributed as follows: WNT (n = 70), SHH (n = 223), Group 3 (n = 144), and Group 4 (n = 326). Expression values were log2-transformed (log2(x + 1)) before downstream analysis.

### Transcriptome Signature Score

To estimate pathway-level activity, signature scores were computed for each sample as the mean log2-transformed expression of predefined gene sets. Gene signatures were defined based on experimentally validated gene sets previously associated with epithelial-to-mesenchymal transition (EMT), ferroptosis, and iron metabolism. EMT-related genes were selected based on established transcriptional signatures and scoring approaches (29,30). Ferroptosis-related genes were selected based on key regulators of lipid peroxidation and ferroptosis sensitivity identified in functional studies (31–33). Genes involved in iron metabolism were selected based on their established roles in cellular iron homeostasis and cancer biology (2,34). The iron metabolism signature included *FTH1, FTL, TFRC, SLC40A1, SLC11A2,* and *IREB2*; the ferroptosis signature included *GPX4, ACSL4, LPCAT3, SLC7A11*, and *AIFM2*; and the EMT signature included *VIM, ZEB1, TWIST1, SNAI1*, and *CDH2*. Differences in signature scores across molecular subgroups were assessed using one-way analysis of variance (ANOVA), followed by Tukey’s HSD post hoc test for multiple comparisons. P-values were adjusted using the Benjamini–Hochberg method. All statistical analyses were performed in R (version 4.4.3; R Foundation for Statistical Computing, Vienna, Austria) using the packages tidyverse, ggplot2, ggpubr, and rstatix. Data visualization was performed using boxplots overlaid with individual data points and colorblind-friendly palettes (Okabe–Ito).

### Gene expression analysis

Gene expression analysis was performed for selected genes score involved in iron metabolism, ferroptosis, and EMT. Differences in gene expression across MB molecular subgroups were first assessed using one-way analysis of variance (ANOVA). When significant, post hoc pairwise comparisons were performed using Tukey’s honestly significant difference (HSD) test to correct for multiple testing. For visualization purposes, pairwise comparisons were additionally displayed using two-sided Student’s t-tests, with p-values reported using standard notation (*p < 0.05, **p < 0.01, ***p < 0.001). Expression distributions were visualized using violin plots overlaid with boxplots and individual data points and colorblind-friendly palettes (Okabe–Ito).

## Statistics analysis

In all figures, data are presented as the mean ± standard error of measurement (SEM) unless otherwise stated in the text. Comparisons between multiple groups were performed using one-way or two-way analysis of variance (ANOVA) followed by Tukey’s post hoc test as appropriate. Survival curves were analyzed using the Kaplan–Meier method with log-rank test.

Statistical analyses were performed using GraphPad Prism (version 10.6.1, GraphPad Software, San Diego, CA, USA). A p-value < 0.05 was considered statistically significant.

## Results

### Genetic deletion of *FTH1* does not impact viability and redox homeostasis of MB cells *in vitro*

To investigate the role of ferritin heavy chain (FTH) in MB, we used two biologically distinct MB models: DAOY cells representing the SHH subgroup with TP53 mutation and HD-MB03 cells representing Group 3. CRISPR-Cas9-mediated *FTH1* deletion was confirmed by immunoblotting (Figure 1A,B). Under basal conditions, FTH loss did not significantly affect viability, intracellular iron levels, ROS production, or lipid hydroperoxide accumulation in either cell line (Figure 1C–J), indicating that MB cells *in vitro* (regardless of MB subgroup) can adapt to ferritin deficiency in the absence of stress.

**Figure 1.**
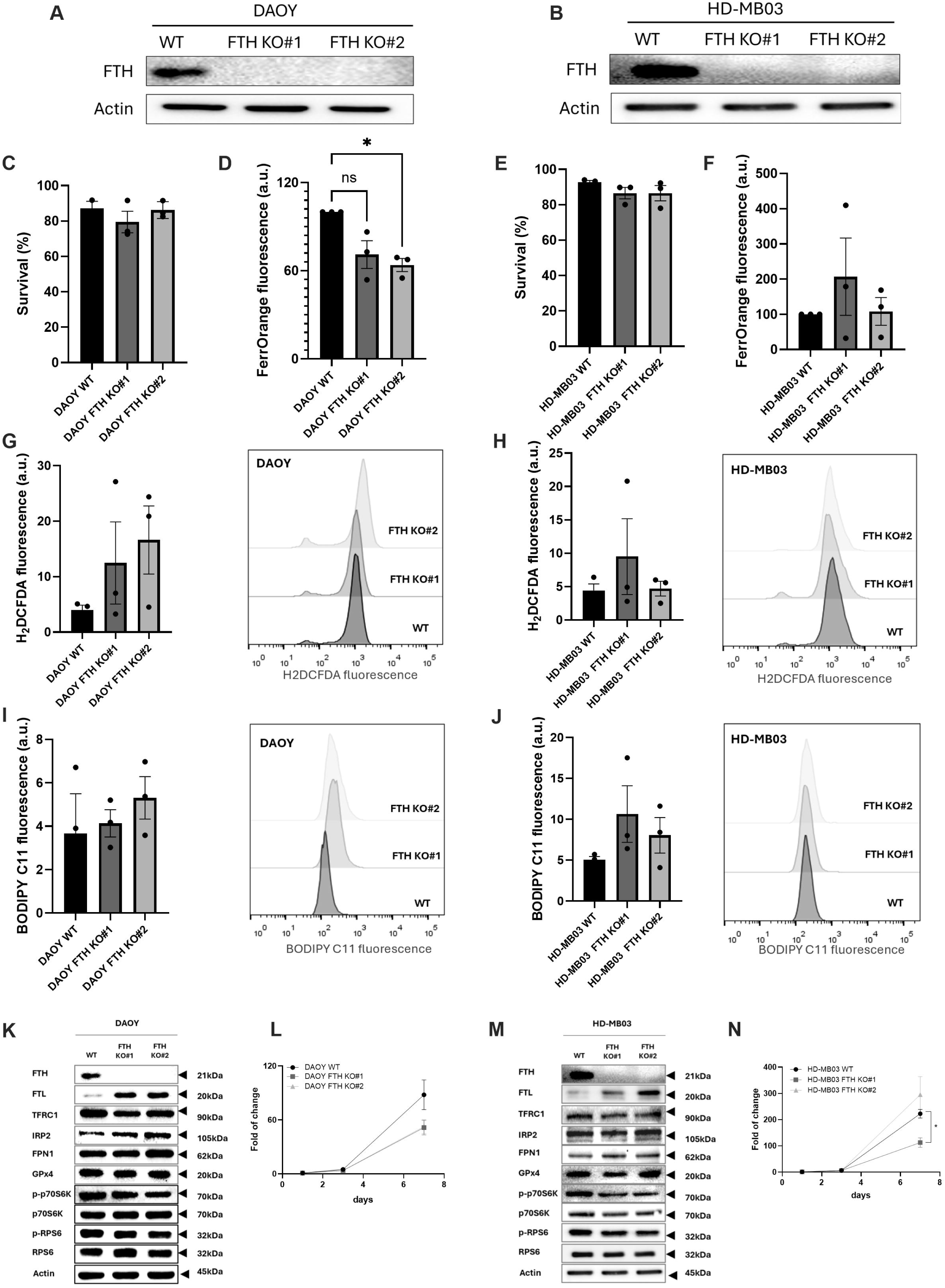
**Deletion of FTH in MB cells does not affect redox homeostasis or survival *in vitro.*** The FTH subunit of the ferritin complex was examined in DAOY (A) and HD-MB03 (B) cells using two independent FTH KO clonal lines (#1 and #2). Basal cell survival was evaluated in DAOY (C) and HD-MB03 (E) WT and FTH KO cells by PI staining followed by flow cytometry analysis. Basal intracellular ferrous iron (Fe²⁺) levels were quantified by using FerroOrange probe in DAOY (D) and HD-MB03 (F) WT and FTH KO cells. Fe²⁺ content was normalized to cell number. To assess a hallmark of ferroptosis, intracellular ROS levels were determined in DAOY (G) and HD-MB03 (H) WT and FTH KO cells. Basal lipid hydroperoxide level was analyzed in DAOY (I) and HD-MB03 (J) WT and FTH KO cells. Molecular adaptations of DAOY (K) and HD-MB03 (M) cells upon FTH deletion were evaluated by immunoblotting. The relative abundance of proteins involved in iron homeostasis (FTH, FTL, IRP2, TFRC1, and FPN1) was determined. mTORC1 pathway activity was assessed through the expression and phosphorylation status of 70S6K and RPS6. GPX4 expression was also analyzed as an anti-ferroptotic marker. Cell proliferation was assessed in DAOY (L) and HD-MB03 (N) WT and FTH KO cells. *Bar graphs represent the mean ± SEM (n = 3). Western blots and histograms are representative of three independent experiments. For WB analysis actin was used as a loading control. Proliferation rates are presented as fold of change (mean ± SEM; n = 3). All analyses were performed 24h post-seeding. *, P < 0.05, comparison with corresponding WT control cells*. *Abbreviations used: Medulloblastoma (MB), ferritin heavy chain (FTH), propidium iodide (PI), reactive oxygen species (ROS), ferritin light chain (FTL), iron regulatory protein 2 (IRP2), transferrin receptor 1 (TFRC1), ferroportin-1 (FPN1), Ribosomal protein S6 kinase (p70S6K), Ribosomal protein S6 (RPS6), glutathione peroxidase 4 (GPx4)*.

To determine how MB cells compensate for ferritin loss, we examined key regulators of iron metabolism (Figure 1K,M; Supplementary Figure 1A,B). Both DAOY and HD-MB03 FTH KO cells upregulated ferritin light chain (FTL), suggesting a shared compensatory attempt to preserve iron storage capacity. However, the adaptive response was markedly stronger in DAOY cells. DAOY FTH KO cells exhibited increased expression of IRP2 and the anti-ferroptotic enzyme GPX4, a trend toward elevated ferroportin (FPN1), reduced TFRC1 expression, and decreased phosphorylation of the mTORC1 downstream targets p70-S6K and RPS6 (Figure 1K; Supplementary Figure 1A). In contrast, HD-MB03 FTH KO cells exhibited only minimal changes in expression of these proteins apart from FTL upregulation and an approximately 50% reduction in IRP2 levels (Figure 1M; Supplementary Figure 1B). These differences likely reflect intrinsic subgroup-specific biology. HD-MB03 cells harbor c-Myc amplification, previously associated with altered ferritin regulation (35,36) which may reduce dependence on ferritin-mediated iron storage, whereas DAOY cells may rely more strongly on ferritin to maintain iron homeostasis and ferroptosis resistance. Consistent with this notion, the two MB cell lines exhibited marked morphological differences (Supplementary Figure 1C), with DAOY cells displaying an elongated, spindle-like morphology, whereas HD-MB03 cells appeared more epithelial-like. This observation was further supported by Western blot analysis showing higher expression of the mesenchymal markers N-cadherin and vimentin, as well as the EMT-associated transcription factors Twist and ZEB1, in DAOY cells compared with HD-MB03 cells (Supplementary Figure 1D).

Given the influence of iron metabolism and mTORC1 signaling on proliferation, we assessed *in-vitro* cell growth following *FTH1* deletion. Although both DAOY and HD-MB03 FTH KO cells displayed reduced proliferation compared with WT controls, statistical significance only occurred in one HD-MB03 clone (Figure 1L,N).

### FTH deletion markedly prolongs survival in orthotopic MB xenografts

To extend our *in vitro* findings, we evaluated the tumorigenic potential of *FTH1*-deficient MB cells in brain orthotopic xenograft models. In a stark contrast to our results from *in vitro* experiments, mice implanted with DAOY FTH KO cells exhibited strikingly prolonged survival rates of approximately three-fold compared to mice bearing WT tumors (Figure 2A). In contrast, *FTH1* deletion in HD-MB03 cells resulted only in a non-significant trend towards improved survival (Figure 2B). Immunohistochemical analysis using Ki67 and nestin confirmed tumor identity and revealed no major differences in proliferative capacity between WT and FTH KO tumors at endpoint (Figure 2A,B). This suggests that ferritin loss primarily affects tumor adaptation and stress tolerance rather than baseline proliferation as was observed *in vitro*.

**Figure 2.**
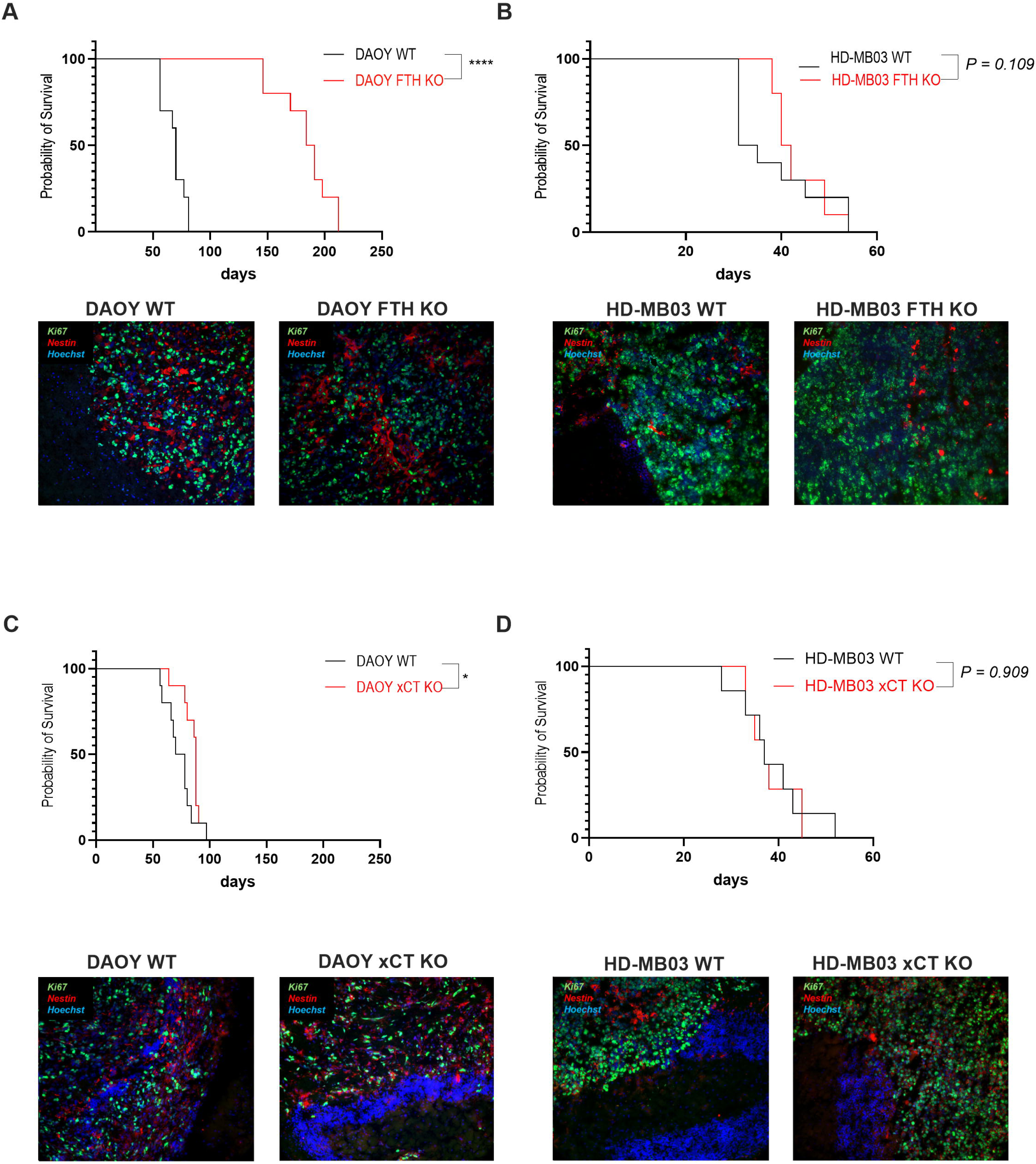
Genetic deletion of FTH reduces tumor growth *in vivo*. Survival curves showing the proportion of surviving mice over time following orthotopic tumor implantation of WT (black lines) and FTH KO (red lines) cells from the DAOY (A) and HD-MB03 (B) cell lines (n = 10). DAOY (C) and HD-MB03 (D) WT and xCT KO clones, were orthotopically implanted into the cerebellum of nude mice. Survival curves are shown for each group (n = 10). Representative images of tumor sections stained with the indicated markers: proliferating cells detected by Ki67 immunofluorescence (green), nestin (neuronal stem cell marker; red), and nuclear DNA counterstained with Hoechst 33342 (blue) are shown below corresponding survival curves. *The images are representative of at least three independent tumors.* *Abbreviations used: ferritin heavy chain (FTH)*

To compare ferritin deficiency with disruption of a canonical ferroptosis defense mechanism, we examined MB cells lacking the cystine/glutamate antiporter xCT (SLC7A11). Previously generated xCT KO cells (22) displayed a substantially stronger ferroptotic phenotype *in vitro* than FTH KO cells, including rapid lipid peroxide accumulation and complete cell death in the absence of alternative cysteine donors (Supplementary Figure 2A–F). xCT deficiency was also associated with amino acid starvation signaling, suppression of mTORC1 activity, and deregulation of iron metabolism (Supplementary Figure 2G). These data were consistent with observations previously reported in pancreatic ductal adenocarcinoma xCT KO models (37). Despite these pronounced *in vitro* effects, xCT deletion only modestly impaired orthotopic tumor growth *in vivo*. DAOY xCT KO tumors showed a statistically significant delay in progression (Figure 2C), but the effect was considerably weaker than that observed following *FTH1* deletion (Figure 2A). In contrast, HD-MB03 xCT KO xenografts exhibited virtually no survival benefit (Figure 2D). Together, these findings suggest that ferritin-mediated iron buffering may play a more critical role in MB tumor adaptation and progression *in vivo* than canonical cystine-dependent ferroptosis defense, despite inducing a comparatively weaker ferroptotic phenotype *in vitro*.

Given the distinct responses of DAOY and HD-MB03 cells to both FTH and xCT deletion, we investigated MB subgroup-specific transcriptional programs related to EMT, ferroptosis, and iron metabolism for potential explanations of these discrepancies. Analysis of patient-derived MB transcriptomic datasets ((28), GSE85217) revealed significant variation in EMT, ferroptosis, and iron metabolism signatures across the four MB molecular subgroups (Supplementary Figure 2H). EMT and iron metabolism signatures were significantly enriched in SHH tumors compared with Group 3 and Group 4 tumors which is consistent with SHH tumors being associated with a more mesenchymal phenotype (Supplementary Figure 1C,D). In contrast, ferroptosis-related signatures were elevated in non-SHH subgroups. These findings support a model in which SHH tumors exist in a more iron-dependent and mesenchymal-like state, whereas Group 3 tumors exist in a more epithelial state and display enhanced activation of ferroptosis-protective pathways, potentially contributing to their reduced sensitivity to FTH loss in our HD-MB03 cell models. Importantly, both *FTH1* and *FTL* expression were substantially lower in Group 3/4 tumors than in WNT/SHH tumors (Supplementary Figure 2I), further supporting a reduced dependence on ferritin-mediated iron storage in non-SHH MB. In contrast, canonical anti-ferroptotic regulators, including *xCT*, *GPX4*, and *FSP1*, did not show similar MB subgroup-specific differences (Supplementary Figure 2I), suggesting that differential ferritin expression may represent a more distinctive feature of SHH versus non-SHH MB than general ferroptosis defense pathways.

### FTH KO cells exhibit increased sensitivity to ferroptosis

Given the central role of ferritin in intracellular iron buffering, we hypothesized that *FTH1*-deficient cells, although adapted to ferritin loss under basal conditions, would be highly sensitive to fluctuations in iron availability. Consistent with this, both DAOY and HD-MB03 FTH KO cells displayed markedly reduced clonogenic survival following long-term exposure to the iron donor ferric ammonium citrate (FAC) compared with WT controls (Figure 3A,B). In DAOY cells, this effect was completely rescued by the ferroptosis inhibitor ferrostatin-1 (Fer-1), whereas HD-MB03 FTH KO cells showed limited response to Fer-1 treatment. In contrast, iron deprivation using the chelator deferoxamine (DFO) affected DAOY WT and FTH KO cells similarly (Supplementary Figure 3A), suggesting that ferritin is more critical for protection against iron excess than as a long-term iron reservoir during depletion.

**Figure 3.**
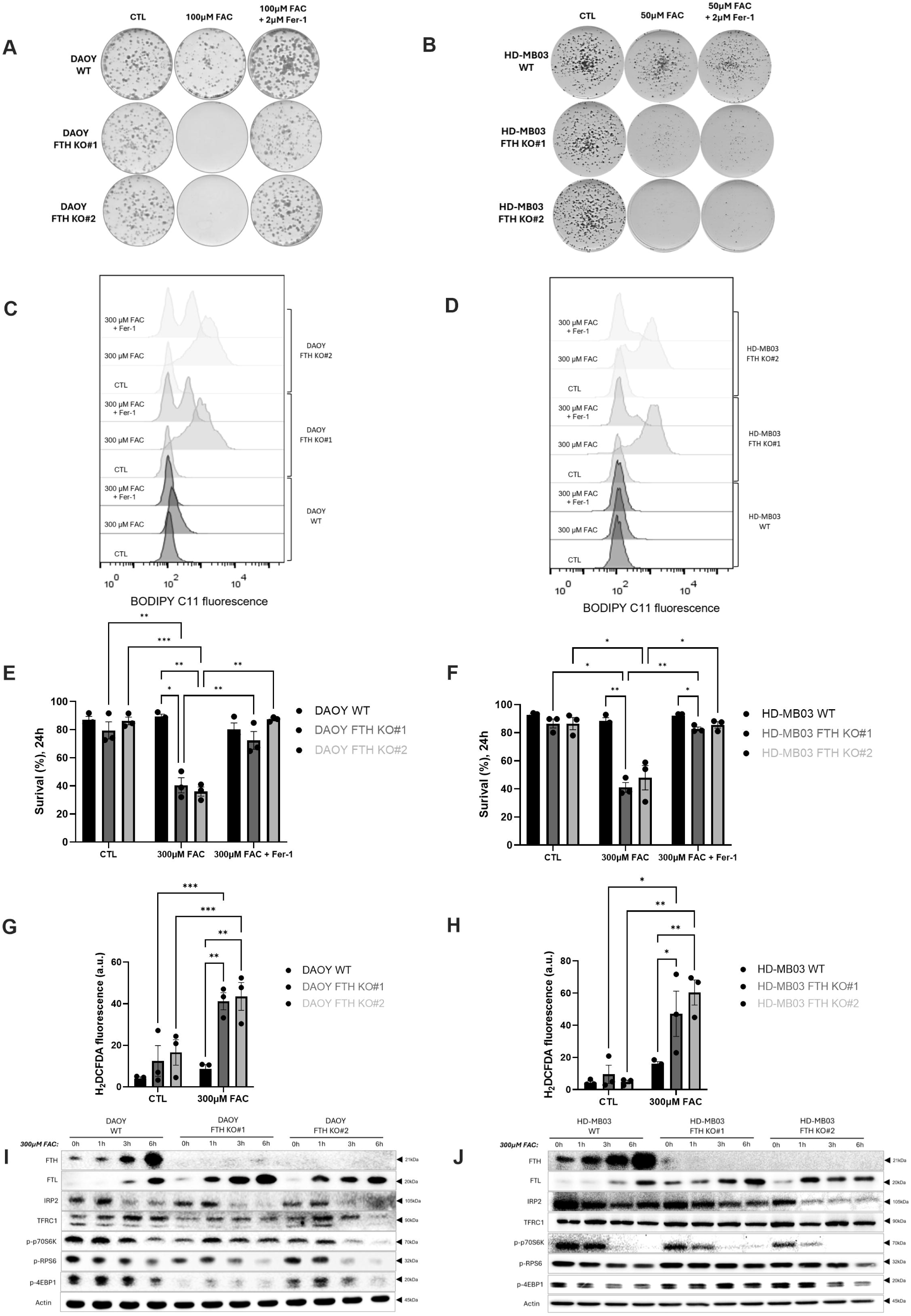
FTH loss sensitizes MB cells to iron-induced oxidative stress and ferroptosis. Seven-day clonal growth of DAOY (A) and HD-MB03 (B) WT and FTH KO cells cultured in media +/- 50–100 µM FAC and 2 µM Fer-1. Lipid hydroperoxide levels (C,D), cell survival (E,F), and ROS production (G,H) were measured in DAOY and HD-MB03 WT and FTH KO cells 6 h or 24 h (cell death) after treatment with 300 µM FAC. Fer-1 was used as a ferroptosis inhibitor. Molecular adaptation of DAOY (I) and HD-MB03 (J) WT and FTH KO cells following treatment with 300 µM FAC for 1 h, 3 h, and 6 h. Relative protein levels of iron metabolism regulators (FTH, FTL, IRP2 and TFRC1) and markers of mTORC1 activity, including phosphorylation of 70S6K, RPS6, and 4EBP1, were evaluated. *Images, blots and histograms are representative of three independent experiments. Bar graphs are presented as mean ± SEM; n = 3. *, P < 0.05; **, P < 0.01; ***, P < 0.001; statistical comparisons are shown between indicated groups*. *Abbreviations used: ferric ammonium citrate (FAC), ferrostatin-1 (Fer-1), reactive oxygen species (ROS), ferritin heavy chain (FTH), ferritin light chain (FTL), iron regulatory protein 2 (IRP2), transferrin receptor 1 (TFRC1), Ribosomal protein S6 kinase (p70S6K), Ribosomal protein S6 (RPS6), eIF4E-binding protein 1 (4EBP1)*.

Mechanistically, short-term FAC exposure induced pronounced lipid hydroperoxide accumulation in FTH KO cells from both models, which was partially rescued by Fer-1 (Figure 3C,D; Supplementary Figure 3B,C). Prolonged FAC treatment triggered robust ferroptotic cell death in both DAOY and HD-MB03 FTH KO cells, which was fully prevented by Fer-1 (Figure 3E,F). This was accompanied by increased ROS levels specifically in FTH KO cells, whereas WT cells remained largely unaffected (Figure 3G,H). Notably, HD-MB03 cells displayed somewhat greater resistance to ferroptosis induction at higher confluency (data not shown). Lipidomics analysis showed that DAOY and HD-MB03 cells enrich polyunsaturated fatty acids in distinct phospholipids, which could contribute to this difference (Supplementary Figure 3D,E), although more investigation is needed to confirm this possibility. In addition, the data suggests that lipid composition is not the major driver for the accumulation of lipid hydroperoxide in FTH KO cells, as we did not observe increased polyunsaturated phospholipids in them. Indeed, polyunsaturated phospholipids tend to decrease in FTH KO DAOY cells, and remain mostly unchanged in FTH KO HD-MB03 cells (Supplementary Figure 3E). Thus, lipid hydroperoxide accumulation and ferroptosis sensitivity in FTH KO cells seemed to be a direct consequence of a default on iron buffering.

To investigate how MB cells adapt to iron overload, we analyzed the dynamics of iron-regulatory proteins following FAC treatment (Figure 3I,J). FAC rapidly induced ferritin expression, with increased FTH levels in WT cells and increased FTL levels in both WT and FTH KO cells observed within 1–3 h. This response correlated with rapid downregulation of IRP2, indicating efficient sensing of elevated intracellular iron. In contrast, TFRC1 expression remained largely unchanged during early time points, suggesting a delayed suppression of iron uptake pathways. These findings support a model in which ferritin induction for iron sequestration represents the primary early defense against an acute iron overload, while modulation of iron import occurs more slowly. Consequently, FTH KO cells likely fail to buffer excess iron before iron uptake pathways can be sufficiently suppressed, ultimately leading to ferroptotic cell death.

Iron overload also strongly inhibited mTORC1 signaling in both MB models, with suppression of p70-S6K and RPS6 phosphorylation evident within 1–3 h of FAC exposure (Figure 3I,J). Although previous studies demonstrated that iron depletion suppresses mTORC1 activity (38–40), our findings indicate that iron excess similarly disrupts mTORC1 signaling, suggesting that this metabolic pathway is highly sensitive to fluctuations in intracellular iron levels either above or below homeostatic levels.

Finally, since FTH KO cells exhibited enhanced susceptibility to iron-induced ferroptosis, we examined their sensitivity to erastin, the canonical inducer of ferroptosis. According to the clonogenicity test, FTH KO cells from both MB models were markedly more sensitive to erastin-induced cell death than WT controls, and this effect was fully reversed by Fer-1 (Supplementary Figure 3F). Together, these findings demonstrate that ferritin plays a critical protective role against ferroptosis independently of the initiating trigger.

### High dose of Vitamin C induces an iron-dependent, but non-ferroptotic cell death in MB cells

Given that FTH KO cells exhibit increased sensitivity to iron overload (Figure 3), we next investigated whether modulation of iron redox state by VitC, without directly increasing total iron levels, would produce a similar effect. Although classically recognized for its antioxidant properties, VitC serves as a natural reducer of iron (41). Specifically, it converts Fe³⁺, including ferritin-bound iron, into Fe²⁺, thereby expanding the labile iron pool (Figure 4A). At pharmacological concentrations, VitC has been reported to induce oxidative stress through redox cycling between Fe²⁺ and Fe³⁺, generating sustained production of hydroxyl radicals that can lead to cell death, frequently described as ferroptosis (42–46). To test this hypothesis, we treated DAOY and HD-MB03 WT cells with a pharmacologically relevant concentration of VitC (10 mM), which is transiently achievable through intravenous or intraperitoneal VitC administration and has been reported to be achievable in multiple clinical trials, without serious side-effects (47–54).

**Figure 4:**
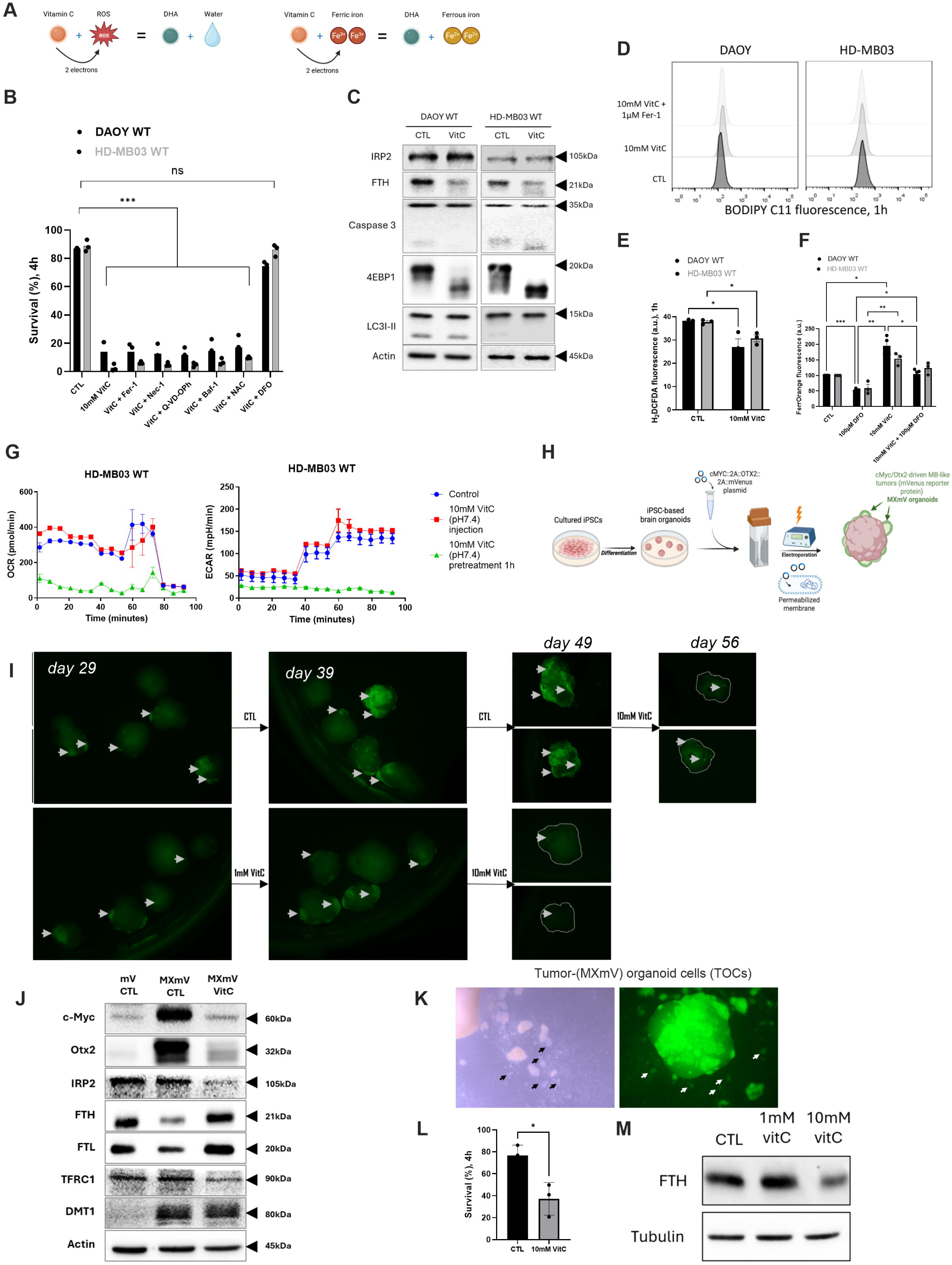
High doses of Vitamin C induce iron-dependent, but ROS-independent cell death distinct from classical ferroptosis. Schematic representation of the redox activity of ascorbic acid (VitC) and its impact on ROS production and Fe³⁺ reduction in the organism (A). Cell viability (B) was assessed in DAOY WT and HD-MB03 WT cells after treatment with a pharmacological dose of VitC (10mM) for 4h in the presence or absence of classical cell death inhibitors, including Fer-1, Nec-1, the pan-caspase inhibitor Q-VD-OPh, Baf-1, antioxidant - NAC or metal chelator - DFO. Molecular responses to VitC treatment were analyzed in DAOY WT and HD-MB03 WT cells after 1 h exposure to VitC (C). Relative protein levels of regulators of iron metabolism (IRP2 and FTH), apoptosis markers (Caspase 3 cleavage), mTORC1 pathway activity (4EBP1) and autophagy marker (LC3) were evaluated by immunoblotting. Lipid hydroperoxide (D), ROS (E), ferrous iron (F) contents, as well as metabolic flux (OCR and ECAR) (G) were measured in DAOY WT and HD-MB03 WT cells after 1 h treatment with VitC. Schematic representation of the generation and maintenance of cerebral organoids from hiPSC (H). The effect of VitC treatment on Otx2/c-Myc-positive tumor cells was evaluated in cerebellar organoids (I). Twenty-nine days post-electroporation with plasmids expressing Otx2/c-Myc and a fluorescent mVenus reporter (MXmV), organoids were treated with low-dose VitC (1mM) for 10 days, followed by high-dose VitC (10mM) for an additional 10 days. A subset of control organoids received only late high-dose VitC treatment. Tumor development was monitored every 4 days by imaging. Relative protein expression was analyzed at the end of the experiments in organoids treated or not with VitC (J). Reduced c-Myc and Otx2 expression was observed in VitC–treated organoids. The abundance of iron metabolism regulators (FTH, FTL, IRP2, TFRC1, DMT1) was also evaluated. Actin was used as loading control. Detached Otx2/c-Myc-positive tumor cells were collected from organoids and used as an independent MB-like cell line, referred to as tumor-organoid cells (TOCs) (K). Cell viability following 4 h treatment with VitC 10 mM was assessed in TOCs (L). FTH protein expression levels were analyzed in response to 1 mM and 10 mM VitC treatment for 1 h (M). Tubulin was used as a loading control. *Images, blots, and histograms are representative of three independent experiments. Bar graphs represent the mean ± SEM; n = 3. *, P < 0.05; **, P < 0.01; ***, P < 0.001; statistical comparisons are shown between indicated groups*. *Abbreviations used: vitamin C (VitC), ferrostatin-1 (Fer-1), necrostatin-1 (Nec-1), pan-caspase inhibitor (Q-VD-OPh), bafilomycin A1 (Baf-1), deferoxamine (DFO), N-acetylcysteine (NAC), ferritin heavy chain (FTH), ferritin light chain (FTL), iron regulatory protein 2 (IRP2), transferrin receptor 1 (TFRC1), divalent metal transporter 1 (DMT1), Ribosomal protein S6 (RPS6), human induced pluripotent stem cells (hiPSCs), tumor-organoid cells (TOCs)*.

VitC induced rapid and extensive cell death in both MB cell lines within 4 h (Figure 4B). Notably, inhibitors of ferroptosis (Fer-1), necroptosis (necrostatin-1, Nec-1), apoptosis (Q- VD-OPh), autophagy (bafilomycin 1, Baf-1), and ROS scavenging (NAC) failed to rescue viability, whereas the iron chelator DFO completely prevented cell death (Figure 4B), indicating that iron toxicity is the primary mediator of VitC-induced cytotoxicity. Consistently, VitC caused a marked reduction in FTH protein levels and strong suppression of mTORC1 signaling, reflected by reduced phosphorylation of 4E-BP1, without evidence of activation of apoptosis (caspase 3) or autophagy (LC3) (Figure 4C).

Unexpectedly, the VitC-induced toxicity differed from canonical ferroptosis. VitC did not elevate lipid hydroperoxides (Figure 4D), despite increased intracellular iron levels and instead reduced ROS levels (Figure 4E,F; Supplementary Figure 4A), explaining the lack of protection by Fer-1 and NAC. Moreover, VitC rapidly induced collapse of both oxidative phosphorylation and glycolysis (Figure 4G), an effect that was not rescued by DFO (Supplementary Figure 4B), suggesting that these conditions disrupted not only free iron homeostasis but also iron-dependent metabolic enzymes.

To extend these findings to a more physiologically relevant model, we used brain organoids in which Otx2 and c-Myc drive Group 3 MB-like tumor formation (MXmV organoids; Figure 4H), as previously described (55,56). Tumor formation became evident approximately 29 days post-transfection (Figure 4I). 10 mM VitC completely eradicated tumors within 10 days, including established tumors treated at later stages, whereas 1 mM VitC had minimal effect (Figure 4I). Tumor burden, monitored by mVenus fluorescence (Figure 4I; Supplementary Figure 4C), correlated with high c-Myc and Otx2 expression, both of which were markedly reduced following VitC treatment (Figure 4J), indicating a selective targeting of MB-like tumor cells.

Interestingly, unlike MB cell lines, organoids responded to VitC with increased FTH and FTL expression alongside reduced IRP2 and TFRC1 levels (Figure 4J). This suggests that activation of adaptive ferritin-mediated buffering occurred in these physiologically relevant 3-dimensional conditions in remaining healthy organoid cells in response to elevated intracellular iron. MXmV transformed cells also displayed invasive behavior characterized by spontaneous tumor cell dispersal (Figure 4K). Tumor organoid-derived cells (TOCs), which expressed Otx2, c-Myc, and mVenus, remained highly sensitive to 10 mM VitC, exhibiting ∼40% increased cell death within 4 h (Figure 4L). Notably, FTH expression in TOCs was dose-dependent, increasing after 1 mM VitC but markedly decreasing following 10 mM treatment (Figure 4M), further supporting ferritin dynamics as a critical determinant of adaptation to iron stress.

Our observations, together with those of other groups, indicate that mesenchymal signatures correlate with increased iron sensitivity ((37,57,58), Supplementary Figure 1C,D; Supplementary Figure 2H,I), led us to examine whether cellular phenotype influences susceptibility to VitC-induced toxicity. Culturing DAOY cells in 3D promoted a more epithelial-like phenotype (Supplementary Figure 4D) and significantly reduced VitC-induced cell death compared with conventional 2D cultures (Supplementary Figure 4E,F). This reduced sensitivity correlated with decreased expression of mesenchymal markers including N-cadherin and vimentin (Supplementary Figure 4E). Interestingly, VitC reduced FTH expression in 2D cultures but increased it in 3D cultures (Supplementary Figure 4E), suggesting that epithelial-like cells possess greater capacity to accommodate iron-associated stress under more physiological-like conditions. Collectively, these findings suggest that high dose of VitC induces a rapid, iron-dependent metabolic collapse in MB through mechanisms distinct from canonical ferroptosis and highlight ferritin dynamics and cellular phenotypes as key determinants of sensitivity to iron-mediated redox perturbation.

### Ferritin-deficient MB cells exhibit increased sensitivity to high doses of VitC

Considering that high-dose VitC (10 mM) reduced FTH expression and rapidly induced iron-dependent cell death, whereas lower sub-lethal concentrations increased ferritin levels in WT cells (Supplementary Figure 5A,B), we investigated whether lower doses of VitC could phenocopy the selective vulnerability of FTH KO cells observed during FAC-induced iron overload (Figure 3).

Time-course analyses demonstrated that lower dose of VitC induced a transient reduction in IRP2 levels in WT cells, occurring after 3h in DAOY cells (Figure 5A) and 1h in HD-MB03 cells (Figure 5B), followed by partial recovery at 6h. TFRC1 expression showed only modest changes, whereas FTH and FTL progressively increased in WT cells, consistent with activation of ferritin-mediated iron buffering. FTL was similarly upregulated in FTH KO cells, which already exhibited elevated basal FTL levels (Figure 5A,B). Importantly, these responses closely resembled the iron-adaptive program induced by FAC treatment (Figure 3I,J), suggesting that VitC-mediated iron reduction is sensed by the cell as a functional iron overload, thereby activating ferritin-dependent homeostatic pathways.

**Figure 5.**
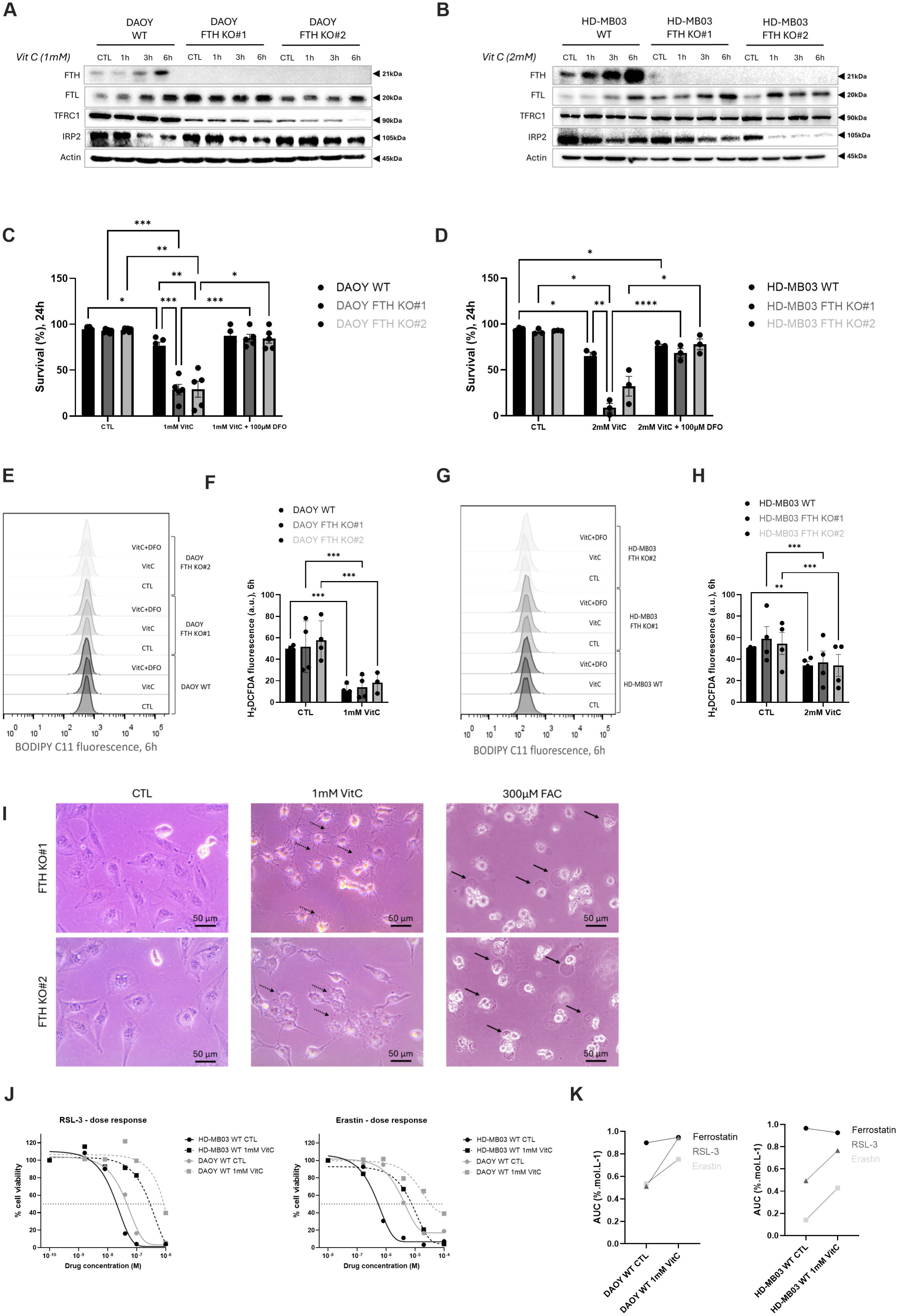
Ferritin protects MB cells against Vitamin C–induced iron toxicity. Expressions of iron metabolism regulators FTH, FTL, TFRC1 and IRP2, were analyzed by immunoblotting in DAOY (A) and HD-MB03 (B) WT and FTH KO cells after short-term treatment with 1 or 2 mM VitC for 1, 3, or 6 h. Cell viability (C,D), lipid hydroperoxide (E,G) and ROS (F,H) contents were assessed in DAOY and HD-MB03 WT and FTH KO cells after 6h or 24 h (cell death) VitC treatment. DFO (100 µM) was used to rescue the observed phenotype in FTH KO cells. (I) Representative micrographs of DAOY FTH KO cells treated or untreated with 1 mM VitC or 300 µM FAC for 24 h. Dotted arrows indicate morphological features of VitC-induced cell death, while solid arrows denote characteristic membrane bubbling observed during ferroptosis. (J) Dose–response curves of DAOY WT or HD-MB03 WT cells treated with erastin or RSL-3 for 72 hours in combination with DMSO (CTL) or VitC 1mM (K) AUC values of ferrostatin, erastin and RSL-3 in DAOY WT and HD-MB03 WT cells either combined with DMSO (CTL) or VitC 1mM. *Histograms and blots are representative of three independent experiments. Bar graphs represent mean ± SEM; n = 3. *, P < 0.05; **, P < 0.01; ***, P < 0.001; statistical comparisons are shown between indicated groups. Abbreviations used: ferritin heavy chain (FTH), ferritin light chain (FTL), iron regulatory protein 2 (IRP2), transferrin receptor 1 (TFRC1), vitamin C (VitC), reactive oxygen species (ROS), deferoxamine (DFO), ferric ammonium citrate (FAC)*.

Functionally, lower dose of VitC caused minimal toxicity in WT DAOY and HD-MB03 cells (Figure 5C,D), indicating effective adaptation to moderate iron redox perturbation. In contrast, FTH KO cells from both models exhibited pronounced sensitivity, with >70% cell death after treatment (Figure 5C,D). This effect was fully rescued by DFO, confirming that cytotoxicity remained strictly iron-dependent. Similar to observations with 10 mM VitC, low-dose treatment did not increase lipid hydroperoxide accumulation (Figure 5E,G) and instead reduced ROS levels in both WT and FTH KO cells (Figure 5F,H), arguing against an initiation of canonical ferroptosis to explain the cytotoxicity. The morphological characteristics observed in VitC-treated cells support the occurrence of a non-ferroptotic cell death mechanism (Figure 5I), in contrast to the morphology exhibited by cells treated with the ferroptosis inducer FAC. Furthermore, 1 mM VitC significantly decreased the sensitivity of MB cells to the canonical ferroptosis inducers erastin and RSL3, while no effect was observed with Fer-1, as assessed in dose response experiments (Figure 5J,K). By using the difference in AUC between the combinatorial treatment and the monotherapy condition, we defined those combinations as antagonistics. Indeed, our data revealed a strong antagonist interaction between VitC and ferroptosis inducers with calculated AUC differences of -22 and -29% mol/L respectively for erastin and RSL3 in WT HD-MB03 and of -22 and -43% mol/L respectively for erastin and RSL3 in WT DAOY (Figure 5K). This is reflected by a strong increase in the IC50 of erastin and RLS-3 when combined with 1 mM VitC (Figure 5J, dotted lines). This effect may be explained by the significantly lower ROS levels observed following VitC treatment (Figure 5F,H), as ROS are required for the Fenton reaction and the subsequent induction of ferroptotic cell death. Together, these findings identify ferritin induction as a critical adaptive mechanism that enables MB cells to tolerate moderate increases in labile iron, whereas MB FTH-deficient cells remain highly vulnerable even to sublethal perturbations in iron redox balance.

To further investigate the role of cellular phenotype in iron sensitivity, we attempted to induce EMT in HD-MB03 cells. Although TGF-β failed to induce detectable changes (data not shown), bFGF treatment produced molecular and morphological features consistent with a partial mesenchymal transition (Supplementary Figure 5C,D). This was accompanied by increased FTH expression, reduced TFRC1 levels, and increased ACSL4 expression, suggesting coordinated remodeling of iron metabolism and membrane lipid composition. Lipidomic analysis revealed that bFGF treatment induced selective remodeling of the phospholipid landscape consistent with a more ferroptosis-prone phenotype. At the bulk level, most major lipid classes remained unchanged, except for a global increase in cholesteryl esters following bFGF exposure (Supplementary Figure 5E). At the phospholipid subclass level, bFGF-treated cells displayed a reduction in plasmalogen phosphatidylethanolamines (PE P-) (Supplementary Figure 5F), a lipid subclass that is seen to have antioxidant properties (59). In contrast, diacyl PE levels were unchanged, but its acyl chain composition was altered (Supplemental Figure 5G,H). We found a trend for reduced abundance of arachidonic acid (20:4)-containing diacyl PE and increased levels of longer-chain polyunsaturated diacyl PE species, particularly 22:4-containing diacyl PE (Supplementary Figure 5G). Notably, enrichment of 22:4 was observed across multiple lipid classes and represented one of the most consistent changes detected by volcano plot analysis (Supplementary Figure 5I). As 22:4 (adrenic acid) is generated through elongation of arachidonic acid, these findings suggest that bFGF promotes PUFA elongation, potentially through activation of ELOVL-dependent pathways. Collectively, the reduction in antioxidant plasmalogen PE species together with the accumulation of highly polyunsaturated phospholipids, including 22:4-containing PE species previously implicated in ferroptosis execution (60), raises the possibility that bFGF induces lipid remodeling in a direction associated with increased ferroptosis sensitivity, similar to that observed in canonical mesenchymal/TGF-driven cellular states.

To determine whether these lipid alterations translate into enhanced vulnerability to iron-dependent oxidative stress, we next assessed the response of FGF-treated cells to high doses of VitC. Importantly, bFGF-pretreated HD-MB03 cells displayed significantly increased sensitivity to VitC treatment compared with untreated controls (Supplementary Figure 5J). Given that pharmacological dose of VitC promotes iron redox cycling, these findings support the notion that FGF-driven lipid remodeling creates a cellular state that is more susceptible to iron-mediated toxicity. Together, these results suggest that acquisition of mesenchymal-like features sensitizes MB cells not only to canonical ferroptosis-inducing conditions but also to broader forms of iron-dependent oxidative stress.

### FTH deficiency confers enhanced sensitivity to iron overload *in vivo*

To assess the *in vivo* sensitivity of FTH-deficient cells to iron overload, we employed the HD-MB03 orthotopic model, which in our previous experiments exhibited only a modest - survival advantage following *FTH1* loss (Figure 2B). Mice bearing either WT or FTH KO tumors were randomized into three groups: a control group, a treatment group receiving the iron donor Fe-dex (a clinically approved formulation used for the management of anemia (61–63)) and a treatment group receiving pharmacological dose of VitC. Fe-dex administration did not confer a significant survival benefit in mice harboring WT tumors. In contrast, mice bearing FTH KO tumors exhibited a statistically significant extension in survival following Fe-dex treatment, indicating increased susceptibility of ferritin-deficient tumors to iron-induced stress *in vivo* (Figure 6A) and recapitulating the enhanced iron sensitivity observed *in vitro* (Figure 3).

**Figure 6.**
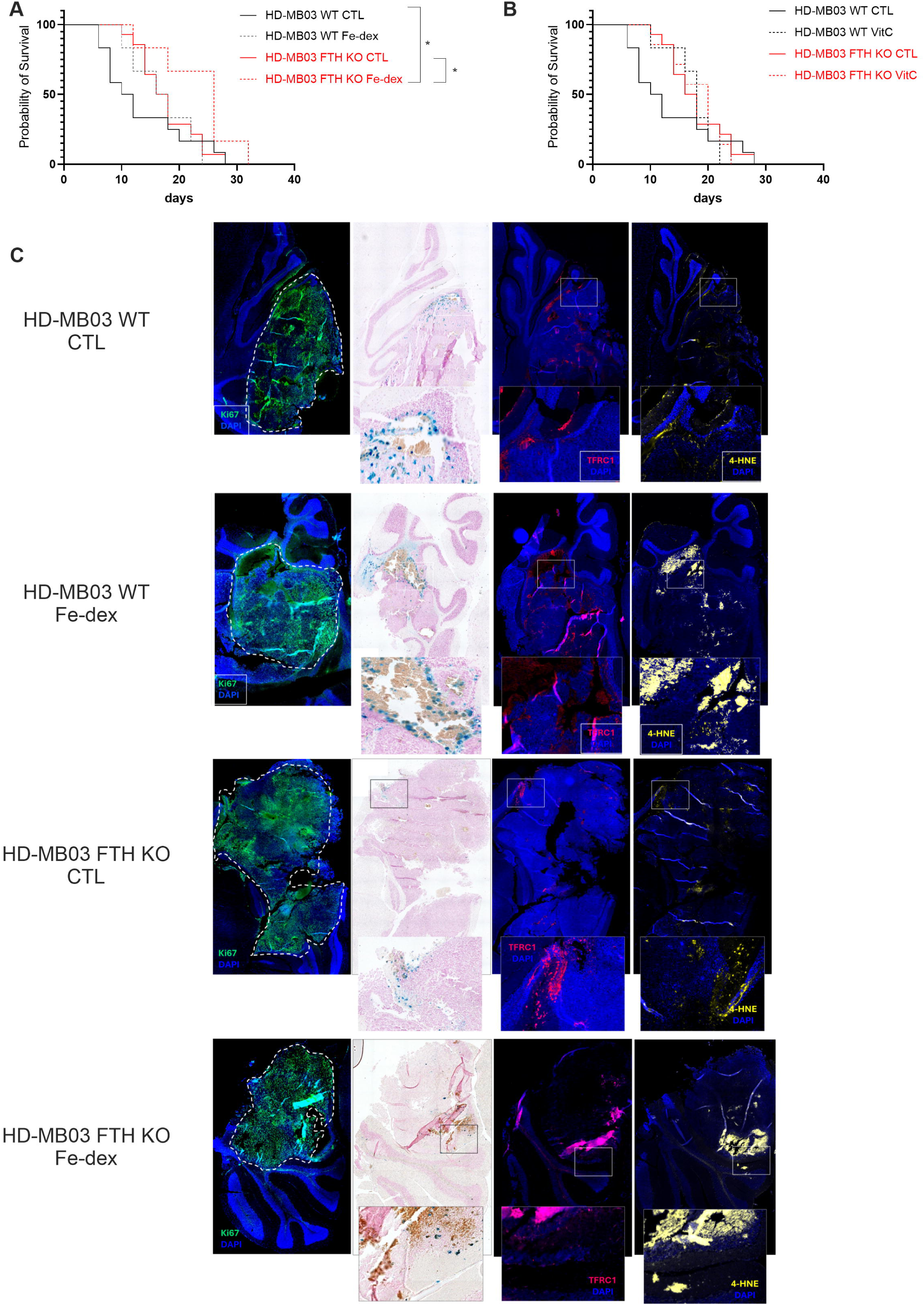
FTH deficiency increases tumor sensitivity to iron overload and ferroptosis *in vivo*. (A) Survival curves (n = 10) showing the proportion of surviving mice over time following orthotopic tumor implantation of HD-MB03 WT (black lines) and FTH KO (red lines) cells treated or not with Fe-dex 500 mg/kg; dashed and solid lines, respectively). Iron dextran was administered once daily for 5 consecutive days by intraperitoneal injection, followed by a second treatment cycle after a one-week treatment-free interval. Time on the graphs is expressed in days starting from the first iron dextran injection. (B) Overall survival of mice bearing orthotopic HD-MB03 WT (black lines) or FTH KO (red lines) tumors following treatment or no treatment with VitC (3 g/kg) (solid versus dashed lines, respectively; n = 10). VitC was administered intraperitoneally twice daily for 2 weeks. (C) Representative images of paraffin-embedded tumor sections of HD-MB03 WT and FTH KO tumors, mice treated or not with Fe-dex. Immunofluorescence staining was performed to detect proliferating cells (Ki67, green), the iron importer TFRC1 (red), the ferroptosis marker 4-HNE (yellow), and nuclear DNA counterstained with DAPI (blue). Corresponding paraffin sections stained for ferrous iron (Fe²⁺) using Prussian blue are also shown. *The images are representative of at least three independent tumors. *, P < 0.05; statistical comparisons are shown between indicated groups.* *Abbreviations used: iron-dextran (Fe-dex), bioluminescence imaging (BLI), ferritin heavy chain (FTH), transferrin receptor 1 (TFRC1), 4-hydroxynonenal (4-HNE), 4′,6-diamidino-2-phenylindole (DAPI)*.

Consistent with the survival findings, whole mount histological iron staining performed on tumor sections at the experimental endpoint revealed substantial accumulation of iron deposits in both Fe-dex-treated and untreated WT tumors, predominantly localized around large blood vessels (Figure 6C, second column). In contrast, iron deposition in FTH KO tumors, irrespective of treatment, was markedly reduced. This observation closely parallels the immunostaining pattern of the principal iron importer TFRC1 within the tumor mass (Figure 6C, third column). Since Prussian blue staining specifically detects Fe³⁺, positive staining therefore primarily reflects iron stored in ferritin, as well as other storage forms present within the tumor tissue. The reduced staining observed in FTH KO tumors therefore supports the conclusion that cellular ferritin constitutes the principal, although not exclusive, component of iron storage and trafficking within the tumor. Residual Prussian blue detected in FTH KO tumors may result from uptake of host-derived ferritin or from alternative intracellular iron storage forms such as hemosiderin, as previously described in glioblastoma (64).

Interestingly, immunostaining for the lipid peroxidation marker 4-HNE was detected in all experimental groups (Figure 6C, fourth column). However, the signal was markedly stronger in both Fe-dex–treated groups, reaching statistical significance only in FTH KO Fe-dex-treated indicating increased oxidative damage associated with iron supplementation (Supplementary Figure 6A). Importantly, this analysis was performed at the experimental endpoint, long after treatment administration, and therefore likely reflects a persistent footprint of oxidative stress rather than the peak damage occurring during iron exposure. Thus, although both genotypes displayed comparable residual levels of lipid peroxidation at endpoint analysis, the acute oxidative insult during treatment was likely substantially greater in FTH KO tumors. This interpretation is consistent with the improved survival of mice bearing FTH KO tumors despite similar endpoint levels of 4-HNE staining.

In contrast to the pronounced *in vitro* activity of high dose of VitC, no significant effect on survival was observed in either WT or FTH KO tumor-bearing mice following systemic VitC administration (Figure 6C). One possible explanation for this discrepancy is the limited penetration of pharmacological VitC across the blood–brain barrier (BBB), which may prevent sufficient intratumoral accumulation to induce iron-dependent toxicity. Consistent with this interpretation, previous studies reporting antitumor effects of VitC in glioblastoma employed intracranial treatment rather than systemic administration (65). Moreover, unlike humans, mice are capable of endogenous VitC synthesis (66), and therefore systemic VitC metabolism, tissue distribution, and baseline antioxidant homeostasis may differ substantially from the human setting, potentially limiting the suitability of this model for evaluating VitC-based therapeutic strategies.

Given the lack of a significant effect of VitC in the orthotopic *in vivo* model, we sought to determine whether this could be attributed to limited VitC penetration across the BBB. To address this question, we repeated the experiment using subcutaneous xenografts generated by injecting HD-MB03 WT and FTH KO cells into nude mice. Consistent with our hypothesis, FTH KO tumors exhibited substantially slower growth than their WT counterparts, indicating that the subcutaneous microenvironment more readily reveals the vulnerability associated with FTH loss than the intracerebellar setting (Supplementary Fig. 6B,C). Moreover, VitC treatment significantly inhibited tumor growth in the subcutaneous model. While WT tumors showed a moderate reduction in growth, the effect was markedly more pronounced in FTH KO tumors, demonstrating enhanced sensitivity to VitC in the absence of FTH (Supplementary Fig. 6B,C). These findings suggest that the lack of therapeutic efficacy observed in the orthotopic model is likely due, at least in part, to limited VitC penetration across the blood-brain-barrier.

## Discussion

Iron is indispensable for tumor growth, yet excess labile iron is inherently cytotoxic. A cancer cell’s ability to balance this paradox remains incompletely understood. Here, we have identified ferritin as a central determinant of iron tolerance in MB and demonstrated that disruption of iron buffering exposes a potent vulnerability to iron-driven metabolic collapse and cell death. Our findings reveal that the protective effects of ferritin for MB cells is not from basal oxidative stress, but specifically from iron–induced toxicity, and that pharmacological VitC exploits this vulnerability by expanding the labile iron pool to lethal levels. Together, these data establish iron toxicity and ferritin as a therapeutically actionable strategy to pursue in aggressive MB.

Contrary to expectations based on ferritin’s antioxidant reputation, the genetic deletion of *FTH1* in MB cells did not perturb basal redox homeostasis, viability, or lipid peroxidation (Figure 1A–J, Supp Figure 1A,B). To the best of our knowledge, this is the first demonstration that MB cells can fully adapt to the loss of the catalytic ferritin subunit under cell culture conditions. Consistent with this adaptation, upregulation of FTL in FTH-deficient cells likely reflects heightened intracellular iron signaling; however, in the absence of the heavy chain’s ferroxidase activity (67), it is insufficient to sustain functional ferritin-mediated iron buffering. Instead, *FTH1* loss drives a broader rewiring of iron-sensing and metabolic signaling networks, particularly in more mesenchymal DAOY cells (Supplementary Figure 1C,D), including activation of IRP2, suppression of iron import, and inhibition of mTORC1 activity (Figure 1K and M). Combined, these changes likely represent compensatory mechanisms to limit intracellular iron flux and preserve redox homeostasis.

Interestingly, although *FTH1* loss did not produce an overt phenotype under *in vitro* culture conditions, mice bearing FTH KO orthotopic xenografts exhibited significantly prolonged survival compared with those implanted with WT tumors (Figure 2A,B). This survival advantage was particularly striking in the DAOY model (the more mesenchymal SHH MB group), where FTH deletion extended survival approximately threefold, whereas in HD-MB03 xenografts (more epithelial group 3 MB) the prolonged survival was not statistically significant. Notably, genetic ablation of *FTH1* proved more effective at delaying orthotopic tumor progression *in vivo* than deletion of xCT (Figure 2C,D), one of the major canonical anti-ferroptotic regulators. This finding was unexpected given the markedly pronounced ferroptotic phenotype displayed by xCT KO cells *in vitro* (Supplementary Figure 2C-F). In contrast, as mentioned above, no observable ferroptotic phenotype following FTH loss occurred *in vitro*. However, the discrepancy between the *in vitro* and *in vivo* consequences of xCT depletion has also been reported in multiple tumor models by both our group and others (37,68–71). Collectively, these observations further emphasize the limitations of conventional *in vitro* systems for studying redox-dependent processes, as supraphysiological oxygen tension and artificial media composition fail to accurately recapitulate the metabolic and oxidative landscape of the tumor microenvironment.

Importantly, tumor growth suppression induced by *xCT* or *FTH1* deletion was consistently more pronounced in DAOY xenografts than in HD-MB03 tumors. This suggests that fundamental MB subtype-specific differences exist in dependence on ferroptosis-related pathways and iron homeostasis mechanisms. These differential responses likely reflect the distinct molecular and metabolic characteristics of the two MB models. In particular, HD-MB03 cells harbor MYC amplification, a defining feature of Group 3 MB and one of the strongest predictors of poor clinical outcome and aggressiveness (35,36). Wu and collaborators have highlighted a close interplay between c-Myc and iron metabolism, with c-Myc regulating iron homeostasis genes, such as *FTH1* and *IRP2*, to maintain elevated intracellular iron levels (72). Therefore, it is reasonable to predict that these modifications may buffer HD-MB03 cells against the effects of FTH loss. In this context, HD-MB03 cells under basal conditions likely only rely marginally on ferritin-mediated iron storage, thus allowing them to survive and proliferate, despite FTH deletion. In contrast, DAOY cells appear to be more dependent on ferritin as an iron buffer, rendering them particularly vulnerable to FTH loss as observed in orthotopic experiments. Transcriptomic analyses of primary MB datasets identified subgroup-specific iron metabolism programs (Supplementary Figure 2H,I). SHH tumors (modeled by DAOY cells) showed elevated EMT and iron metabolism signatures, whereas Group 3 tumors (modeled by HD-MB03 cells) exhibited lower *FTH1/FTL* expression and higher ferroptosis-related signatures. These findings suggest that MYC-driven Group 3 MB are less dependent on ferritin-mediated iron storage, potentially explaining their reduced sensitivity to FTH loss. Future studies should determine whether EMT status predicts responsiveness to iron-targeted therapies.

Acute ferritin degradation may produce stronger antitumor effects than genetic FTH loss, as shown in patient-derived glioblastoma models where lysosomal delivery of FTH siRNA caused marked redox and metabolic disruption (73). However, pharmacological targeting ferritin remains underexplored. MMRi62, the only reported ferritin-degrading agent, induces ferroptosis-like cell death in pancreatic cancer models, although its mechanism and tumor specificity remain poorly defined (74). The differences between acute FTH inhibition and genetic knockout in our study further suggest that FTH-KO cells acquire adaptive resistance, highlighting potential mechanisms of resistance to future ferritin-targeted therapies.

Although MB cells can survive and proliferate under basal conditions without ferritin-mediated iron storage, these adaptative mechanisms proved insufficient when cells were challenged with excess iron. These data revealed an essential role for ferritin in protecting tumor cells from iron-induced ferroptotic collapse (Figure 3). Notably, HD-MB03 cells, which were otherwise highly resistant to FTH deletion *in vivo*, remained sensitive to ferroptosis as confirmed by treatment with the iron donor Fe-dex significantly extending survival in xenograft experiments and marked increase in 4-HNE staining (Figure 6A,B; Supplementary Figure 6A). Furthermore, our findings reveal a fundamental role for ferritin depletion in a ROS-independent form of iron cellular toxicity, distinct from classical ferroptosis. While oral VitC is limited to ∼200 µM, intravenous administration achieves sustained millimolar levels with demonstrated antitumor activity across multiple cancer types, including brain tumors (45,47,49,50,53,54,75,76). Consistent with clinical glioblastoma observations, 10 mM VitC rapidly induced MB cell death in our current study, an effect rescued solely by DFO which implicates iron mobilization as the key driver of cellular toxicity (Figure 4). VitC caused rapid ferritin depletion with concomitant intracellular iron accumulation independent of lysosomal ferritinophagy (Figure 4B-F), supporting direct ascorbate-mediated iron release (77–82). In contrast to ferroptosis models of cell death (42,43,45,65), VitC reduced intracellular ROS, did not trigger lipid peroxidation, and was not rescued by ferrostatin-1 or NAC, establishing a ROS-independent iron toxicity pathway. Furthermore, and surprisingly, VitC treatment reduced the sensitivity of MB cells to the canonical ferroptosis inducers erastin and RSL3, most likely due to decreased ROS levels required for Fenton chemistry. Early metabolic collapse occurred (Figure 4G) but was not found to be a causative factor for toxicity, as DFO restored viability without correcting metabolism. Importantly, VitC selectively eradicated MB-like tumors in brain organoids while sparing non-transformed tissue, reflecting differential uptake mediated by cancer-associated sodium-vitamin C (SVCT) and glucose transporters (83–85). Ferritin further modulated cytotoxicity in this context with *FTH1*-deficient cells exhibiting hypersensitivity to VitC, with concentrations tolerated by WT cells, resulting in complete elimination of FTH KO cells. These results identify ferritin-mediated iron sequestration as a key determinant of resistance to VitC–induced toxicity.

Based on these defining features, we propose that this distinct form of iron-dependent cell death be termed *ferrioptosis*. *Ferrioptosis* is characterized by rapid ferritin depletion, intracellular mobilization of redox-active iron through ferric-to-ferrous reduction, profound metabolic collapse, and execution of cell death. Specifically, the name reflects the reduction of **ferri**c (Fe³⁺) to **ferro**us (Fe²⁺) iron, emphasizing that iron reduction itself, rather than ROS production through Fenton chemistry, is the critical determinant of cytotoxicity. Importantly, despite their distinct mechanisms of execution, both ferroptosis (12,37,57,58) and ferrioptosis (Supplementary Figure 4 and 5) appear to share a common determinant of sensitivity, with mesenchymal cellular identity conferring increased susceptibility to each form of iron-dependent cell death. Instead of replacing ferroptosis, *ferrioptosis* complements it by extending the conceptual framework of iron toxicity to include iron-dependent cell death mechanisms that do not rely on ROS accumulation or lipid peroxidation. We propose *ferrioptosis* as a conceptual framework rather than a definitive classification, anticipating that future studies will determine whether this mechanism extends beyond pharmacological VitC and potentially to analogous forms of transition-metal toxicity.

Interestingly, although VitC had no significant effect on tumor growth in the orthotopic model, it significantly delayed the growth of both WT and FTH KO tumors in the subcutaneous model, with greater efficacy in FTH KO tumors (Supplementary Figure 6C,D). These results indicate that limited VitC penetration across the blood–brain barrier may underlie the lack of therapeutic effect in the orthotopic setting, rather than an intrinsic resistance of FTH KO cells to VitC *in vivo*. This interpretation is consistent with previous glioblastoma studies that demonstrated antitumor efficacy occurred following intracranial rather than systemic VitC delivery (65). Together, these findings indicate that ferritin’s principal function in MB is to establish the upper threshold of tolerance to iron toxicity, defining a critical safeguard against iron-mediated cell death.

By revealing the full spectrum of cellular liabilities that emerge when the collapse of iron buffering occurs via ferritin removal, our work re-frames iron toxicity as a therapeutically actionable vulnerability, driven not only by oxidative stress and ferroptosis, but also by ROS-independent iron cytotoxicity, aka *ferrioptosis*. Importantly, the adaptive responses uncovered in ferritin-deficient cells provide a mechanistic blueprint for anticipating resistance to ferritin-targeted pharmacological interventions. Together, these findings establish a conceptual and mechanistic foundation for therapeutic strategies that exploit iron toxicity to destabilize tumor homeostasis, transforming an essential nutrient into a selective and potent anticancer liability.

## Acknowledgments

The authors are sincerely grateful to all the associations, donors, and individuals whose generous financial contributions and continued encouragement made this work possible. We particularly thank the Fondation Flavien, Un Nouvel Espoir; the Fédération Enfants Cancers Santé; the Groupement des Entreprises Monégasques dans la Lutte contre le Cancer (GEMLUC); and the Savchuk Foundation for their longstanding and unwavering support. The authors would also like to acknowledge the HiTS drug screening platform of CRCM and in particular Dr Xavier Morelli and Carine Derviaux.

## Author Approvals

All authors have read and approved the final version of the manuscript for submission to bioRxiv. This manuscript has not been published or accepted for publication elsewhere.

## Statement of significance

Medulloblastoma co-opts primitive neurodevelopmental programs that sustain iron-dependent growth. Our findings identify ferritin as the determinant of iron tolerance, establishing a paradigm shift in therapeutic strategy from restricting iron availability to exploiting iron toxicity.

