## Supplementary Figures for "Deletion of Ferritin Heavy Chain Limits Tumor Growth and Promotes Iron-Dependent Stress in Medulloblastoma"

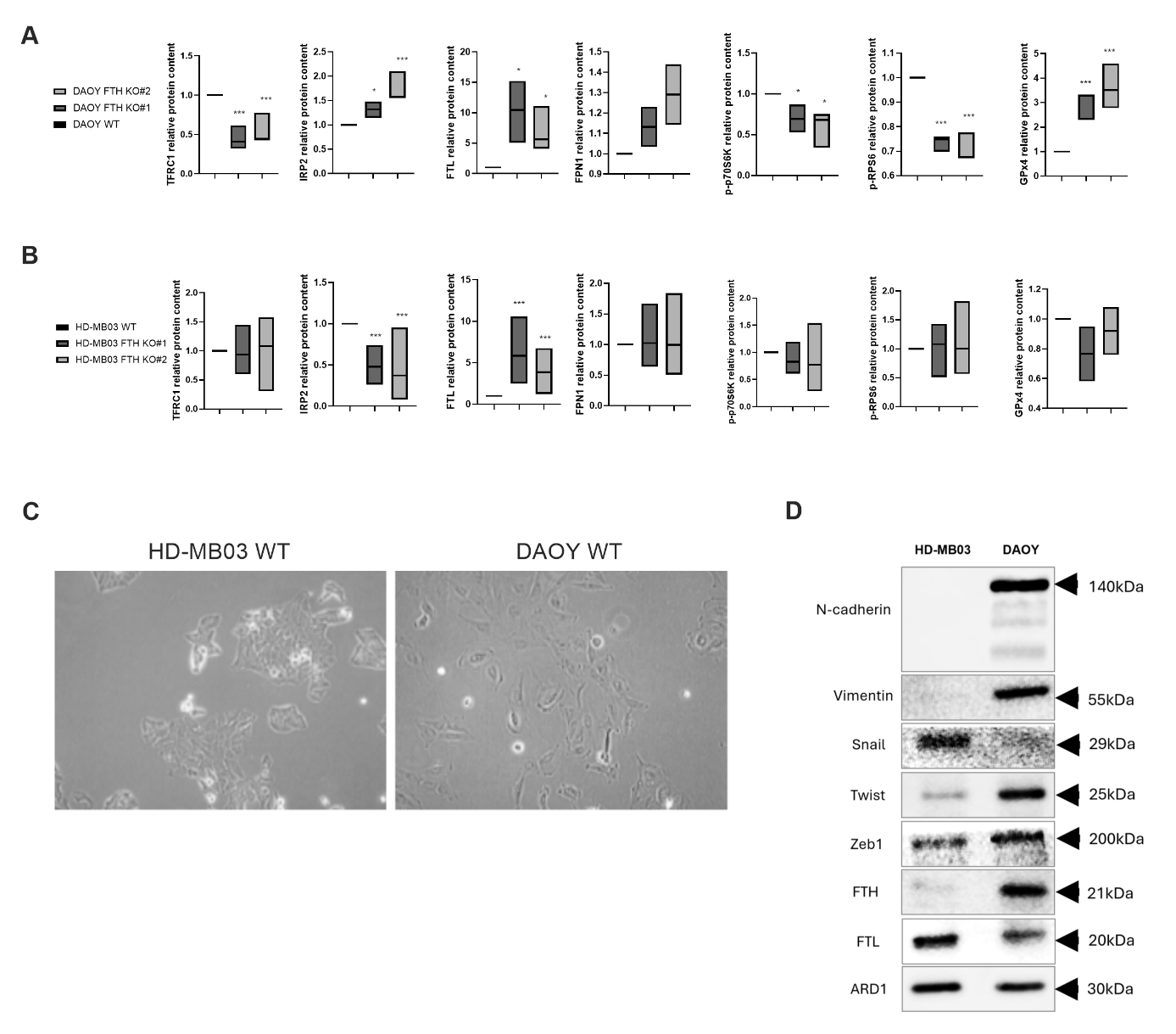
**Supplementary Figures:**

**Supplemented Figure 1** (related to Figure 1)**:**

Quantitative analysis of immunoblot experiments comparing WT and FTH KO DAOY (A) and HD-MB03 (B) cells. See Figure 1K and 1M for corresponding immunoblots.

(C) Representative images illustrating morphological differences between HD-MB03 and DAOY WT cells, with HD-MB03 cells displaying a more epithelial-like phenotype and DAOY cells exhibiting a more mesenchymal-like phenotype.

(D) Basal molecular differences between HD-MB03 and DAOY cells assessed by immunoblotting. Expression levels of epithelial-to-mesenchymal transition (EMT)-associated markers, including N-cadherin, vimentin, Snail, Twist, and Zeb1, as well as ferritin subunits (FTH and FTL), were compared between the two cell lines. ARD1 was used as a loading control.

*Graphs represent mean ± SEM; n = 3. *, P < 0.05; **, P < 0.01; ***, P < 0.001; (A, B) comparison with corresponding WT group; (D,E) statistical comparisons are shown between indicated groups.*

*Abbreviations used: transferrin receptor 1 (TFRC1), iron regulatory protein 2 (IRP2), ferritin light chain (FTL), ferroportin-1 (FPN1), Ribosomal protein S6 kinase (p70S6K), Ribosomal protein S6 (RPS6), glutathione peroxidase 4 (GPx4)*

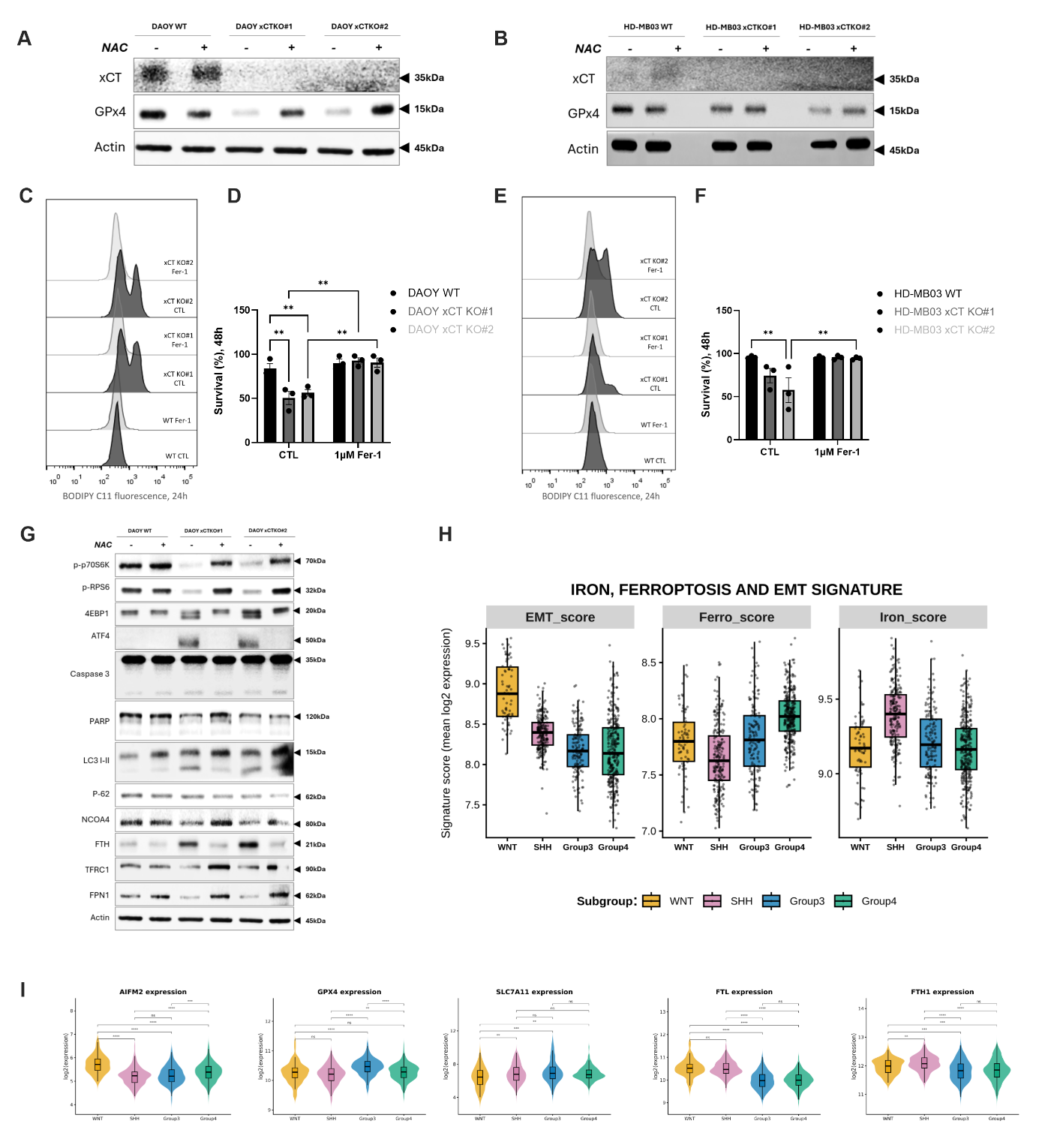

**Supplemented Figure 2**  (related to Figure 2)**:**

Immunoblot analysis of DAOY (A) and HD-MB03 (B) WT and xCT KO cells. Relative protein levels of xCT and GPX4 were evaluated in cells cultured with or without NAC. Actin was used as a loading control.

Lipid hydroperoxide accumulation was measured in DAOY (C) and HD-MB03 (E) WT and xCT KO for 24h with or without 1 µM Fer-1.

Cell viability was assessed following lipid hydroperoxide measurement in DAOY (D) and HD-MB03 (F) WT and xCT KO cells cultured after 48h with 1 µM Fer-1 used to rescue ferroptotic cell death.

Western blot analysis of DAOY WT and xCT KO clones (#1 and #2) was performed culture with or without NAC, to assess mTORC1 activity (p-p70S6K, p-RPS6, and 4EBP1), amino acid stress (ATF4), apoptosis (caspase 3, PARP), autophagy (LC3I-II, p62, NCOA4), iron metabolism (TFRC1, FTH, FPN1) (G)). Actin was used as a loading control.

(H) Transcriptompic analysis of iron metabolism, ferroptosis, and EMT gene signature scores across MB subgroups. Signature scores were calculated as the mean log2-transformed expression of predefined gene sets. Each dot represents an individual tumor sample; boxplots indicate the median and interquartile range. Statistical differences across subgroups (WNT, SHH, Group 3, Group 4) were assessed by one-way ANOVA, followed by Tukey’s HSD post hoc test.

(I) Differential expression of ferroptosis and iron metabolism–related genes across MB subgroups. Violin plots show log2-transformed expression levels of representative genes *AIFM2, GPX4, SLC7A11, FTL, FTH1*, across molecular subgroups (WNT, SHH, Group 3, Group 4). Boxplots indicate the median and interquartile range. Statistical differences were assessed by one-way ANOVA followed by Tukey’s HSD post hoc test.

*Images, blots, and histograms are representative of at least three independent experiments. Bar graphs represent mean ± SEM; n = 3. *, P < 0.05; **, P < 0.01; ***, P < 0.001. Statistical comparisons are shown between indicated groups.*

*Abbreviations used: glutathione peroxidase 4 (GPX4), N-acetylcysteine (NAC), ferrostatin-1 (Fer-1), Ribosomal protein S6 (RPS6), ribosomal protein S6 kinase (p70S6K), ribosomal protein S6 (RPS6), eIF4E-binding protein 1 (4EBP1), activating transcription factor 4 (ATF4), poly (ADP-ribose) polymerase (PARP), microtubule-associated protein 1A/1B-light chain 3 (LC3I-II), transferrin receptor 1 (TFRC1), ferritin heavy chain (FTH), ferroportin-1 (FPN1), epithelial-to-mesenchymal transition (EMT).*

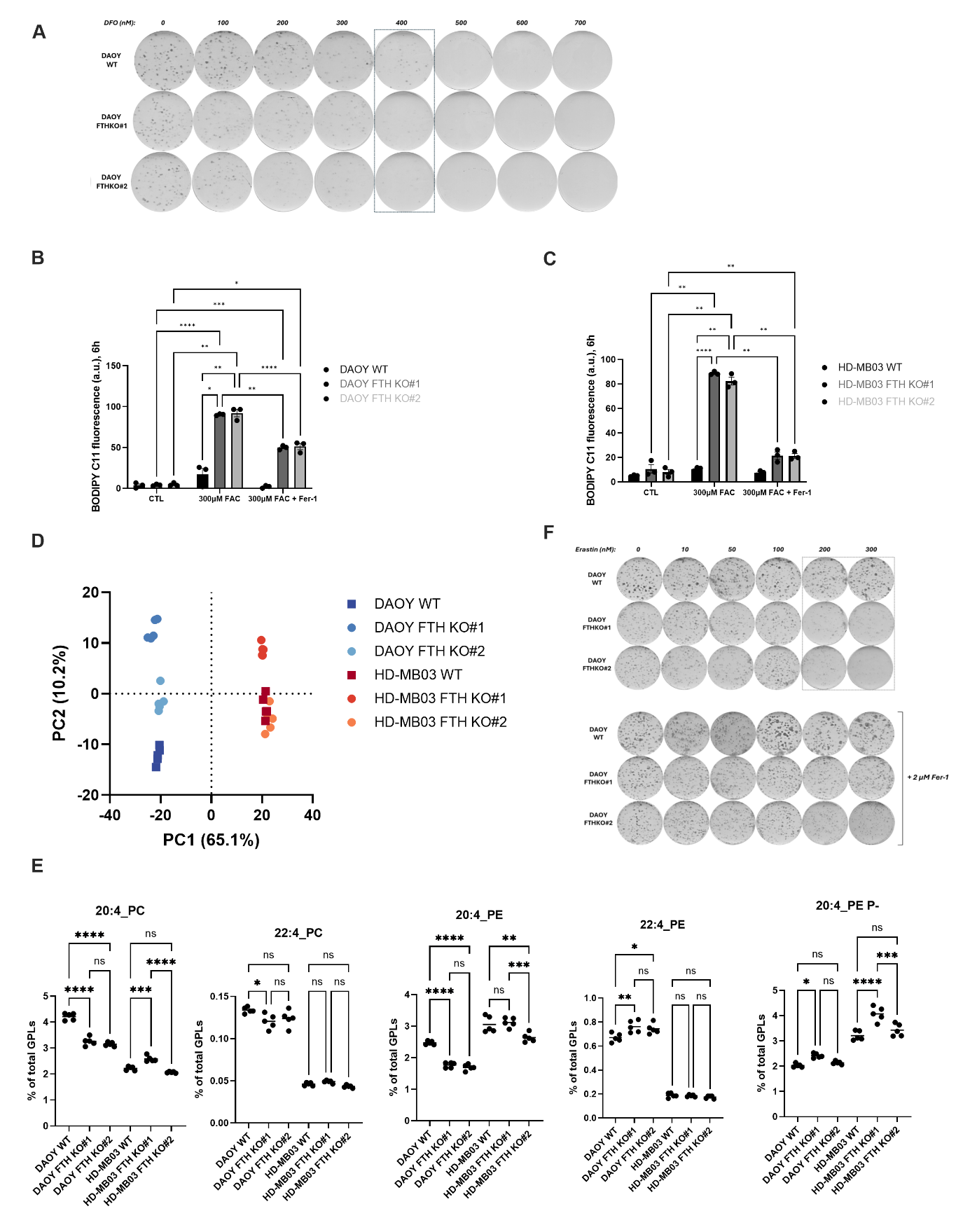

**Supplemented Figure 3**  (related to Figure 3)**:**

(A) DAOY WT cells were cultured in DMEM supplemented with increasing concentrations of DFO (0–700 nM). After 10 days, colonies were visualized by Giemsa staining.

Relative lipid hydroperoxide levels were measured in DAOY (B) and HD-MB03 (C) WT and FTH KO cells following 6 h treatment with 300 µM FAC, with or without 2 µM Fer-1.

(D) Principal component analysis of lipid profiles in the indicated samples. Each point illustrates a biological replicate, and the first two principal component scores are used for plotting.

(E) Comparison of polyunsaturated lipid levels between cell lines and clones. All lipids from the indicated subclass (sharing head groups and sn-1 linkages) were summed when containing the indicated polyunsaturated acyl chain.

(F) Clonogenic growth of DAOY WT and FTH KO clones #1 and #2 was assessed after 7 days of culture in the presence of increasing concentrations of the xCT inhibitor erastin (0, 10, 50, 100, 200, 300 nM), with or without 2 µM ferrostatin-1. Colonies were visualized by Giemsa staining.

*Histograms and images are representative of three independent experiments.*

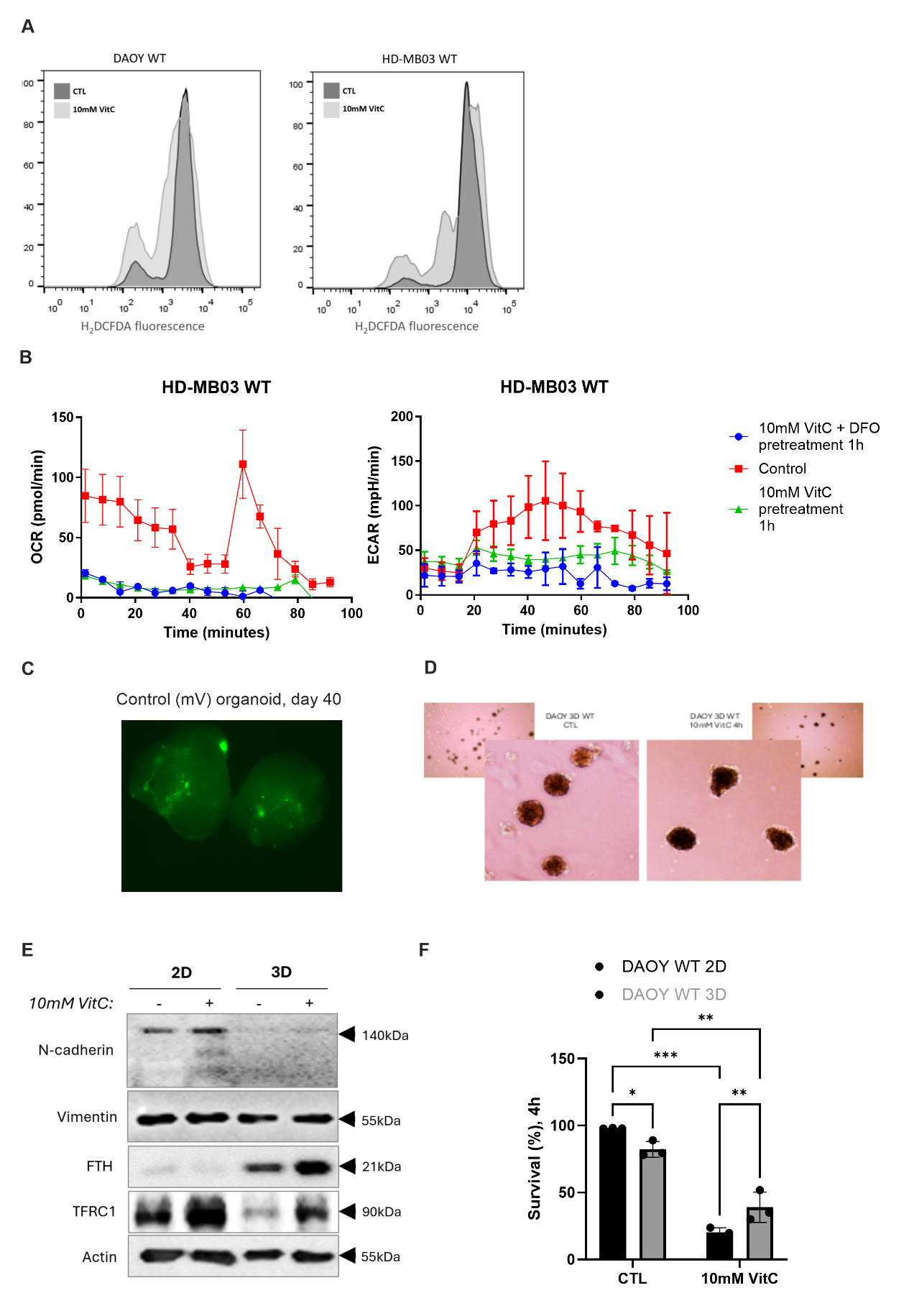

**Supplemented Figure 4**  (related to Figure 4)**:**

(A) Representative histograms of H_2_DCFDA fluorescence in DAOY WT and HD-MB WT cells after treatment with 10 mM VitC (refer to Figure 4E). (B) OCR and ECAR were measured in HD-MB03 WT cells pretreated with 10 mM VitC with or without 100 µM DFO for 1h (B). Data are presented as mean ± SEM; n = 3. (C) Representative image of control (mV) organoids at day 40. Organoids were electroporated with a plasmid expressing only the mVenus reporter gene. (D) DAOY WT cells were seeded in low-adherent plates to form spheres. Treatment with 10 mM VitC induced a bubbling phenotype around spheres, reflecting VitC–mediated cytotoxicity. (E) EMT marker expression (N-cadherin, Vimentin) was compared between 2D and 3D DAOY cells after VitC treatment, along with FTH and TFRC1 as indicators of iron metabolism. (F) Cell viability of DAOY WT cells grown in spheres was measured after 4 h VitC treatment using PI staining, comparing DAOY WT cells in 2D versus 3D culture.

*Histograms, graphs, blots and images are representative of three independent experiments.*

*Abbreviations used: 2′,7′-dichlorodihydrofluorescein diacetate (H_2_DCFDA), vitamin C (VitC), oxygen consumption rate (OCR), extracellular acidification rate (ECAR), deferoxamine (DFO), epithelial-to-mesenchymal transition (EMT).*

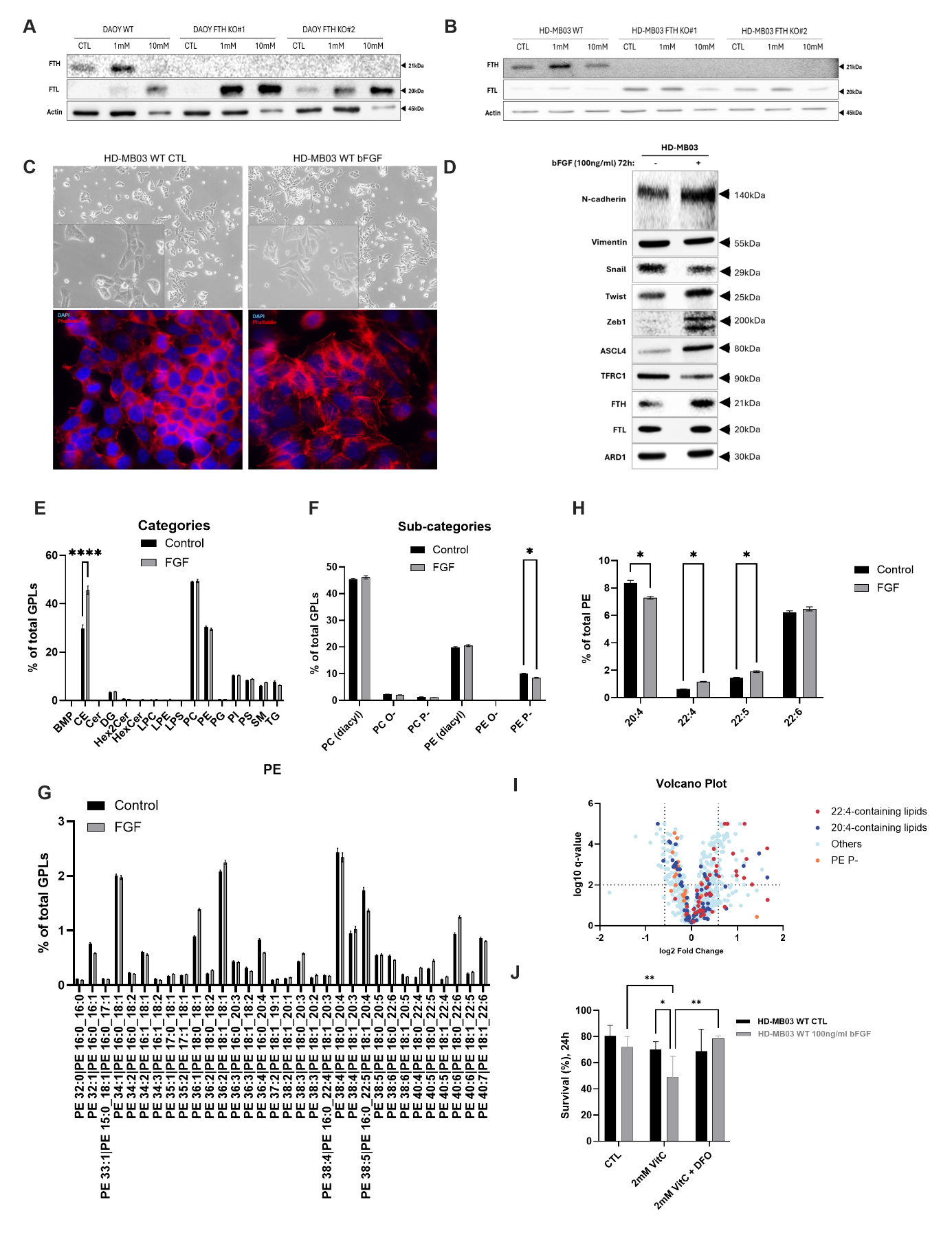

**Supplemented Figure 5** (related to Figure 5)**:**

Relative protein expression of the ferritin subunits FTH and FTL was analyzed in DAOY (A) and in HD-MB03 (B) WT and FTH KO clones #1 and #2. Cells were seeded in media for 24 h and then treated with either 1 mM or 10 mM VitC for 1 h.

HD-MB03 cells, which exhibit a more epithelial phenotype, were cultured with 100 ng/mL bFGF for 72 h. Representative images show the induction of mesenchymal morphology in HD-MB03 cells treated with bFGF, and phalloidin staining highlights cytoskeletal changes under the same conditions (C). Relative protein expression of EMT markers, iron metabolism, and ACSL4 was assessed (D).

(E) Lipid changes associated with bFGF treatment. The sums of lipids from the same lipid classes are shown.

(F) Same as (E), but with distinction of lipid subclasses having distinct sn-1 linkages.

(G) Changes in phosphatidylethanolamine (PE) profiles associated with bFGF treatment.

(H) Analysis of PE acyl chain changes associated with bFGF treatment. The values indicated the contribution of each acyl chain on the total pool of PE acyl chains.

(I) Volcano plot of lipid changes associated with bFGF treatment. Each point illustrates a lipid, which is colored based on the indicated structural feature.

(J) Cell viability of HD-MB03 WT cells pretreated with or without bFGF for 72 h and subsequently exposed to 1 mM VitC for 24 h (H). DFO was used as rescue agents.

*Histograms, images and blots are representative of three independent experiments.*

*Abbreviations used: vitamin C (VitC), epithelial-to-mesenchymal transition (EMT), ferritin heavy chain (FTH), ferritin light chain (FTL), transferrin receptor 1 (TFRC1), acyl-CoA synthetase long chain family member 4 (ACSL4), vitamin C (VitC), basic fibroblast growth factor (bFGF), ARD1 (arrest-defective-1 Protein), deferoxamine (DFO).*

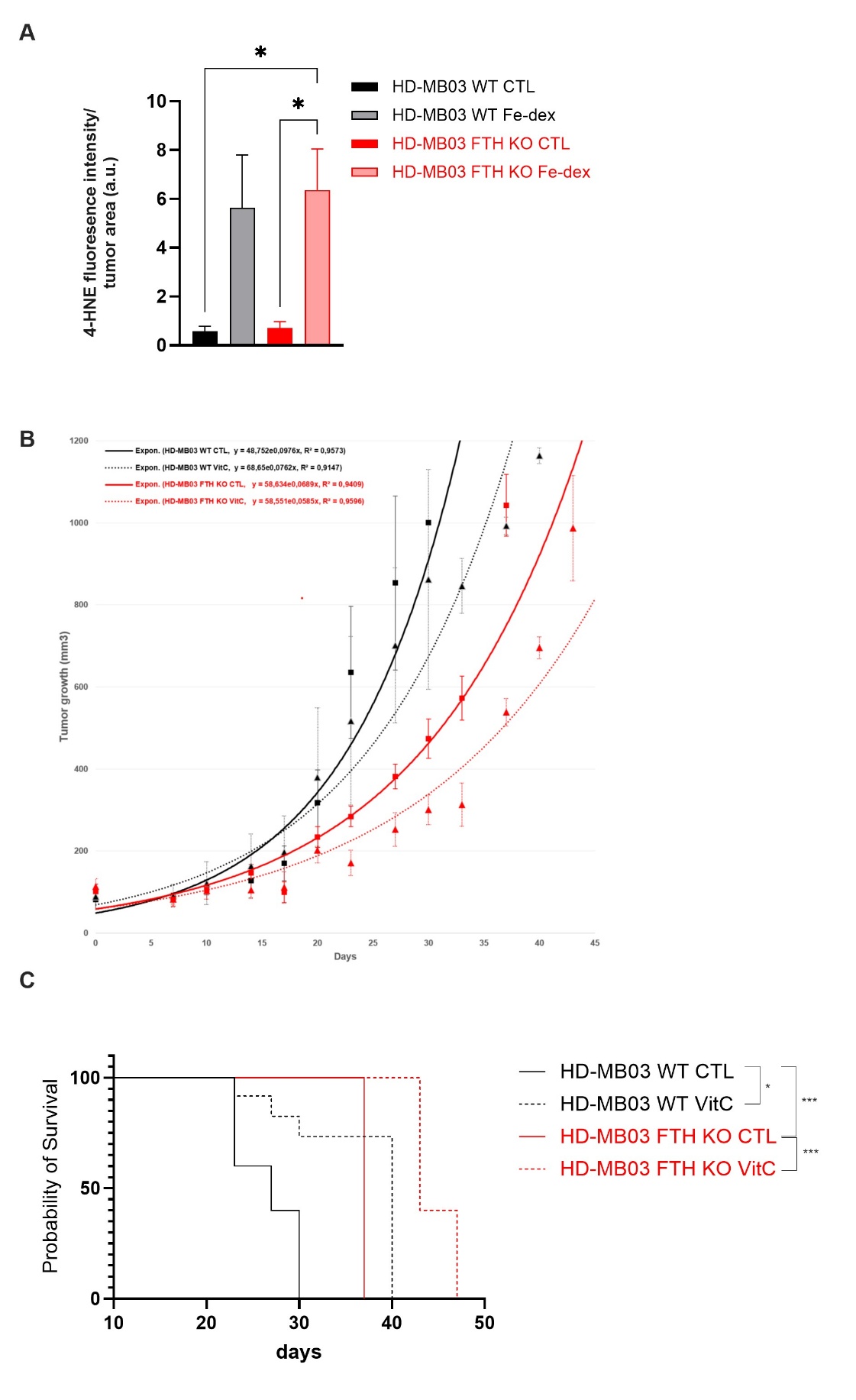

**Supplementary Figure 6 (**related to Figure 6**)**

(A) Quantification of 4-HNE staining in HD-MB03 WT and FTH KO tumors collected from mice treated with or without Fe-dex at the study endpoint (orthotopic model, refer to Figure 6A).

(B) Tumor growth curves of HD-MB03 WT (black lines) and FTH KO (red lines) cells implanted subcutaneously (1 000 000 cells per tumor) into nude mice treated (dashed lines) or untreated (solid lines) with VitC (3 g/kg), administered intraperitoneally twice daily (n = 5).

(C) Overall survival of mice bearing subcutaneous HD-MB03 WT (black lines) or FTH KO (red lines) tumors following treatment with VitC (3 g/kg) (dashed lines) or no treatment (solid lines) (n = 5).

*Bar graphs represent mean ± SEM; n = 3, Tumor growth curves represent average of 5 independent tumors. *, P < 0.05; ***, P < 0.001; statistical comparisons are shown between indicated groups.*

*Abbreviations used: iron-dextran (Fe-dex), vitamin C (VitC), ferritin heavy chain (FTH), 4-hydroxynonenal (4-HNE).*
